# Sector search strategies for odor trail tracking

**DOI:** 10.1101/2021.03.03.433838

**Authors:** Gautam Reddy, Boris I. Shraiman, Massimo Vergassola

## Abstract

Terrestrial animals such as ants, mice and dogs often use surface-bound scent trails to establish navigation routes or to find food and mates, yet their tracking strategies are poorly understood. Tracking behavior features zig-zagging paths with animals often staying in close contact with the trail. Upon sustained loss of contact, animals execute a characteristic sequence of sweeping “casts” – wide oscillations with increasing amplitude. Here, we provide a unified description of trail-tracking behavior by introducing an optimization framework where animals search in the angular sector defined by their estimate of the trail’s heading and its uncertainty. *In silico* experiments using reinforcement learning based on this hypothesis recapitulate experimentally observed tracking patterns. We show that search geometry imposes limits on the tracking speed, and quantify its dependence on trail statistics and memory of past contacts. By formulating trail-tracking as a Bellman-type sequential optimization problem, we quantify the basic geometric elements of optimal sector search strategy, effectively explaining why and when casting is necessary. We propose a set of experiments to infer how tracking animals acquire, integrate and respond to past information on the tracked trail. More generally, we define navigational strategies relevant for animals and bio-mimetic robots, and formulate trail-tracking as a novel behavioral paradigm for learning, memory and planning.

---

Experimental studies demonstrate the ability of ants, dogs, humans, and rodents to track odor trails^1–6^. Rodents accurately track trails in the dark, remaining close to the trail and casting when contact is lost (Figure 1a)^5^. Carpenter ants closely follow a trail while sampling it using a “criss-cross” pattern with their two antennae (Figure 1b)^1^. Current models of this behavior rely on variants of chemotaxis^7^ based on continuous estimates of the rising and falling odor gradients as the trail is crossed. One such strategy compares simultaneous odor concentrations detected by two spatially separated sensors^8^. Yet, rats with a blocked nostril^5^ and ants with a single antenna^1^ are still able to track trails, although less accurately. An alternative chemotaxis strategy has the animal measuring odor gradients along its trajectory and turning when a significant decrease is perceived^5^.

**Figure 1.**
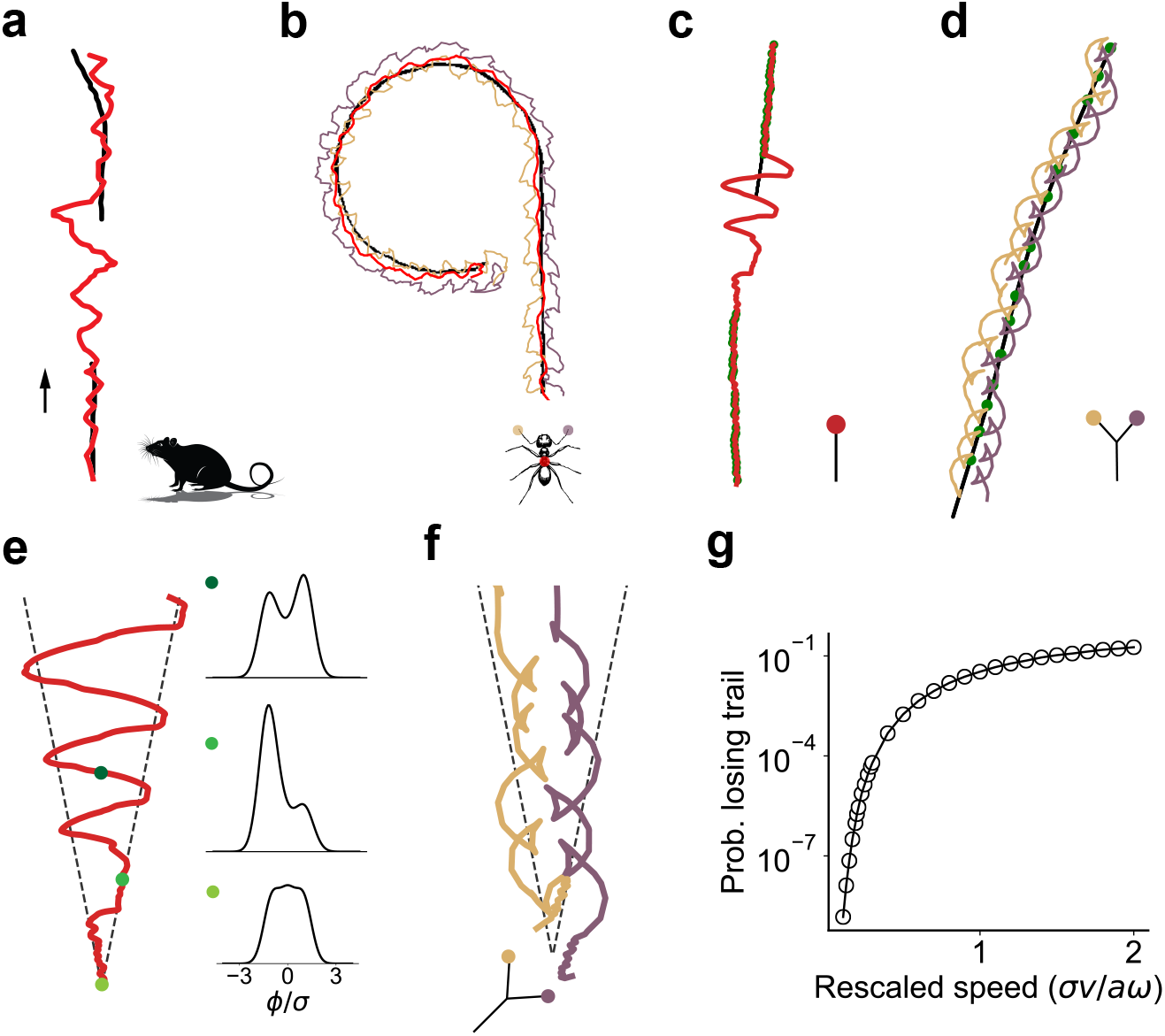
Sample trail tracking trajectories from previous experiments and our reinforcement learning (RL) simulations. (a) A rat (head position in red) tracking a trail (in black). Adapted from ref. 5. Note the wide casts on extended loss of contact with the trail. (b) A carpenter ant tracking an odor trail (black) using a stereotyped “criss-crossing” strategy^1^. (c,d) Sample trajectories obtained from RL for agents with one sensor (panel c) and two sensors (panel d) recapitulate experimentally observed tracking patterns in panels a,b. (e) Left: Search paths executed by RL agents with a single sensor upon loss of contact with the trail. Right: The initial prior distribution (bottom plot) over trail headings transforms into a bimodal posterior distribution (top two plots), which drives the oscillatory pattern of casting. (f) RL agents with two sensors show a characteristic criss-crossing pattern close to the last detection point. The search path is similar to the single-sensor agent at large distances (Figure M1a). (g) RL agents show a trade-off between tracking speed (re-scaled by the sector angle *a*, sensor size a and sampling frequency *ω*) and the probability of losing the trail entirely.

While chemotaxis-based strategies can allow for trail tracking when trails are continuous, they fail when trails are broken and gradients are absent, which is certainly relevant for animals tracking trails in the wild. In experiments with broken trails^1, 5^, the absence of signal triggers casting, which is a fundamental feature shared with olfactory searches in a turbulent medium^9–11^. Even though turbulent searches also feature sporadic cues, airborne odor signals tend to be localized in a cone and, even within the cone, the signal is highly fluctuating^12, 13^. Therefore, beyond qualitative similarities between terrestrial trail tracking and airborne olfactory searches, the specific statistics of detections, geometric constraints and behavioral patterns are distinct.

In contrast with chemotaxis-based algorithms, we propose an alternative framework built on the searcher exploiting past contacts with the trail to maintain an estimate of the trail’s local heading and its uncertainty, which defines an angular sector of probable trail headings that radiates from the most recent detection point. The resulting “sector search” provides a quantitative description of trail-tracking behavior that unifies its various phases and yields specific experimental predictions.

We first show that reinforcement learning (RL) based on the sector search idea can recapitulate natural behavior. An RL agent in this scheme learns to traverse the trail as quickly as possible while minimizing the probability of losing it (see Methods for details). Our *in silico* RL experiments show that general aspects of animal tracking behavior naturally emerge (see Figure 1c,d). Specifically, casts are observed around the most likely heading of the trail, and their amplitude is within the angular sector defined by the initial uncertainty σ of the trail’s heading *ϕ*. The reason for the oscillatory pattern of casting is intuitive. Indeed, while moving along a path 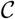 without detecting the trail, the estimated heading’s probability distribution *P*(*ϕ*) (see Figure 1e) is updated into 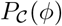 as

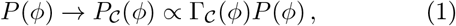

where 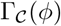 is the probability of not detecting the trail headed along *ϕ*. Irrespective of the explicit form of 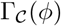, the depletion of headings already explored generally leads to a bimodal posterior distribution, with the two modes at the edges of the angular sector (see Figure 1e). Oscillations are then understood in terms of marginal value theory^14, 15^: we show using a minimal model of casting (Methods) that the turning point of a cast occurs when the marginal value of continuing on one side of the sector is outweighed by the probability of finding the trail on the opposite side.

We proceed now by establishing geometric limits on tracking speed. A typical RL curve for the probability of losing the trail versus speed is shown in Figure 1g. Its monotonicity epitomizes universal limits that “staying on the trail” imposes on tracking speed. Intuitively, searching slowly reduces the distance between detections (the inter-detection interval (IDI)), decreases the uncertainty in the estimate of the trail’s heading, and thus the probability of losing the trail. However, these benefits come at the cost of slow forward progression along the trail. In contrast, moving quickly reduces the detection rate, leading to longer IDIs, increased uncertainty and loss probability.

We quantify the above trade-off using simple scaling arguments. Suppose the tracking agent has a sensor of size *a*, samples at a frequency *ω* and moves with a fixed forward speed v. As shown in Figure 2a, the angle subtended by the detector at distance *r* from the last contact is *a/r*. The agent searching over an angular sector scans then at a rate *dϕ*/*dt* = *ωa*/*r*. Integrating the above expression for the angular rate, we obtain the typical time for re-establishing contact with the trail: *t_c_* ~ *ω*^−1^*e*^*σv*/*aω*^. The corresponding distance *L* ~ *vt*_*c*_ is obtained using *r* ~ *vt*. The heading of the trail is known with uncertainty *σ*, which is the opening angle of the conical sector shown in Figure 2a. Uncertainty is expected to depend on the distance *L′* from the *previous* detection as *σ*(*L′*) = (*L′*/*ℓ*)^*γ*^ where *ℓ* and *γ*, characterize the statistics of trails (see below and Figure 2d). Importantly, a stable strategy for long-term tracking requires that successive IDIs should *on average* be equal, i.e., *L* = *L′*. Combining *L* = *vt_c_* with *L′* = *σ*^1/*γ*^*ℓ* and the expression for *t_c_*, we finally obtain an upper bound on the tracking speed *v*:

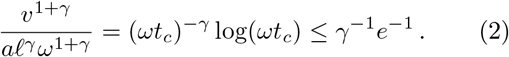

**Figure 2.**
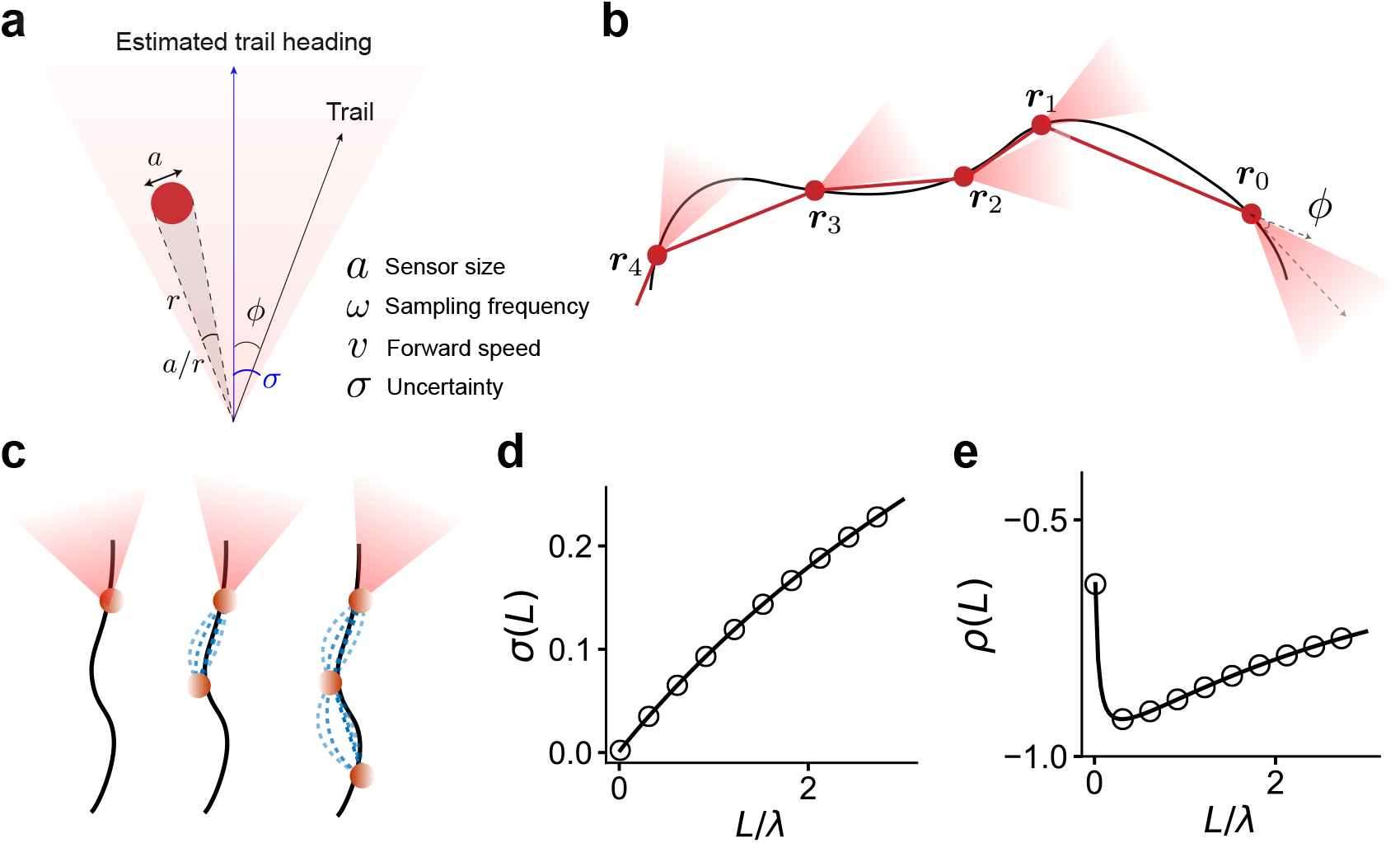
History-dependence and trail models. (a,b) Trail tracking naturally splits into distinct episodes punctuated by trail detections by the searcher. In each episode, we propose that the tracker searches for the trail using an estimate of the trail’s heading updated based on the past points of contact with the trail and a model of trail statistics. We affix a polar coordinate system with the origin at the most recent contact point and the azimuthal angle defined relative to the estimated trail heading. The uncertainty *σ* fixes the angular width of the search. The searcher moves forward with a speed *v* while sampling at a frequency *ω*. A sensor of size *a* spans *a/r* radians at distance r, which determines the rate at which the angular space is searched. (c) To estimate where the trail is headed and its uncertainty from past contacts, the tracker can either use local anisotropy estimated from a single contact (left) or extrapolate from previous points of contact using a model of trail statistics (middle, right). In the latter case, the most likely trail paths (dashed blue lines) are similar to interpolated splines, which capture basic geometric notions of persistence in heading and curvature. (d) The uncertainty in trail heading (in radians) as a function of the distance between points of contact for the GWLC model of trails discussed in the text. (e) The correlation between trail heading at the most recent and second-most recent point of contact for the GWLC model changes with the distance between these points yet it is generally expected to be negative.

Its maximum 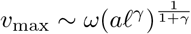 defines the optimal stable tracking speed in terms of the tracker’s sensory parameters and trail statistics. The basic element that leads to this bound is the geometric factor 1/*r* that underlies searching over an angular sector. The result from that *ωt*_c_ is of order one (*e*^1/*γ*^) explains experimental observations (Figure 1) that tracking animals typically take only a few samples to re-establish contact with the trail.

Ideas from polymer physics suggest a statistical description of trails. We ask how detecting the trail at a set of points ***r***_0_, ***r*_1_**, ***r***_2_,…(Figure 2b) constrains its future heading? We consider the case when the searcher keeps track of the two most recent points of contact; a more extended memory is discussed further below. Intuition for the two-point case is provided by the familiar “curve” tool in graphical design software, which draws a cubic spline through a set of prescribed points (Figure 2c). The tool captures the simple intuition that tangents to a curve are continuous, i.e., the trail’s heading has local persistence, which is a plausible, minimal assumption about trails. We show in the SI that cubic spline interpolation corresponds to the most likely path (through a fixed set of points) in the so-called worm-like chain (WLC) ensemble (originally introduced for polymers)^16, 17^. In this ensemble, the tangent direction undergoes diffusion with rate *κ*, and the uncertainty is then 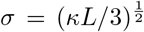, which determines the two parameters *γ*, = 1/2 and *ℓ* = 3*κ*^−1^ in Eq. (2). Actual trails could be smoother and have a well-defined curvature (the rate of change of heading) that persists on a characteristic length scale *λ*. We capture this ensemble of curves by an extended WLC model (EWLC) with two parameters: persistence length *λ* and typical radius of curvature ξ (Methods). Uncertainty is then given by *σ* ≤ *L*/2ξ (hence *γ*, = 1 and ℓ = 2ξ) at distances *L* < *λ* while diffusive scaling is recovered at larger distances with an effective diffusivity *k* = 2*λ*ξ^−2^. Finally, the two models are combined in the general WLC (GWLC) ensemble with crossovers across the various regimes (Methods). In summary, each model leads to a “propagator” which encodes how information about past contacts is integrated to form an estimate of the trail’s heading while taking into account geometric aspects of trails. A feature shared across models is that the headings at two consecutive contacts are anti-correlated (Figure 2e), which reflects the bending of the spline relative to the chord seen in Figure 2c.

Why and when do searchers need to cast? The question stems from our previous result that a few samples are typically sufficient to re-establish contact with the trail. To address it quantitatively, we consider again the setup of Eqs. (1), (2). The non-detection probability averaged over the ensemble of trails that pass through past contact points is

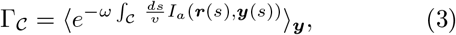

where *s* parametrizes the searcher’s path 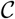 and the Boolean indicator function *I_a_* measures if the agent at ***r***(*s*) is within sensing range *a* of the trail at ***y***, i.e, the integral is the time spent in contact with the trail. Numerical simulations of the search show a power-law scaling regime for 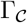, which is cut-off at short distances by the initial surge and at large distances by trails escaping out of the casting envelope (Figure 3a-c). We proceed to explain these three regimes shown in Figure 3b. Intuitively, at short radial distances *r* < *a*/*σ* ~ *v*/*ω* (the latter from Eq. (2)), the sensor covers the entire sector of likely headings, the searcher can just move forward and 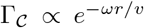 (Figure 3b). Casting sets in if the searcher reaches, without detection, a distance *r* ≳ *a*/*σ*, i.e., when the sector is not fully covered any longer by the sensor of size *a*. The sector geometry in Figure 3a implies that the length of a single casting sweep is proportional to *r*. A fixed forward speed then implies that the distance between successive encounters with the trail also scales with *r*. Hence, the number of times the searcher crosses the trail (and thus the time spent on the trail) per unit radial distance decreases as 1/*r*. This 1/*r* scaling in the overlap then leads to a logarithmic integral in the exponent of Eq. (3) and thus a power-law regime 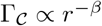 during casting. The optimal exponent β* depends on the statistics of the trails, yet it generally satisfies β* > 1 (β* = 1.63 for the EWLC model, see Eq. (M15)). Since 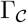 is a cumulative distribution, β > 1 implies that the mean distance is indeed determined by the lower cutoff, i.e., the trail is typically found in a few samples, as estimated in Eq. (2). However, the power-law decay implies that casting phases are frequent and can span up to the upper cutoff where all aspects of trail statistics and search geometry come into play, as discussed next.

How should an agent perform sustained casts so as to minimize the probability of losing the trail while maximizing tracking speed? To go beyond the above scaling arguments, we now consider the geometry of sector search in detail. The searcher’s path is parameterized by the sequence of turning points of its casting trajectory, {*r_k_*, *θ*_*k*_} (in polar coordinates w.r.t the most recent point of contact), which are to be optimized. We maximize the average tracking speed *L/T*, where *L* and *T* are the distance and duration between the most recent and the subsequent point of contact with the trail. As discussed previously, the uncertainty estimated for a bout of sector search depends on the inter-detection interval (IDI). To constrain the uncertainty, we therefore constrain the average, 〈*L*〉. Hence, we consider the following optimization problem:

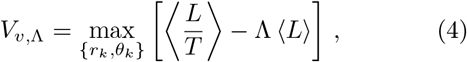

where Λ is the Lagrange multiplier enforcing the 〈*L*〉 constraint. The turning points and the searcher’s speed *v* affect the probability of detecting the trail in a single cast, which is implicit in the expectation in Eq. (4). We solve the Bellman equation corresponding to the above optimization problem using dynamic programming (see Methods), which sequentially optimizes for the turning points by considering at each step the two possibilities that either the trail is detected during the cast or the agent advances to the next cast. The probabilities for these two events are controlled by the non-detection probability given by Eq. (3). In the event of no detection, the estimate of the trail’s heading is updated according to Eq. (1). The resulting optimization yields a search strategy with an increasing sequence of casting angles (Figure 3d). The specific casting strategy depends on how trails meander and curve (Figure 3e). The choice of *v* controls the trade-off between the tracking speed and the probability of losing the trail entirely (Figure 3f). Independent of the choice of *v* and 〈*L*〉, azimuthal excursions are by and large conical but extend dramatically (Figure 3d,e) when trails that were initially inside the cone escape from it with high probability (Figure 3c). This happens at a distance that scales with ℓ but also depends on the sector envelope (which in turn depends on *σ*). Intuitively, at length-scales ~ ℓ the trail’s heading decorrelates from its initial value, the relevance of information on past detections has expired, the trail is effectively lost and it is optimal to stop progressing forward.

**Figure 3.**
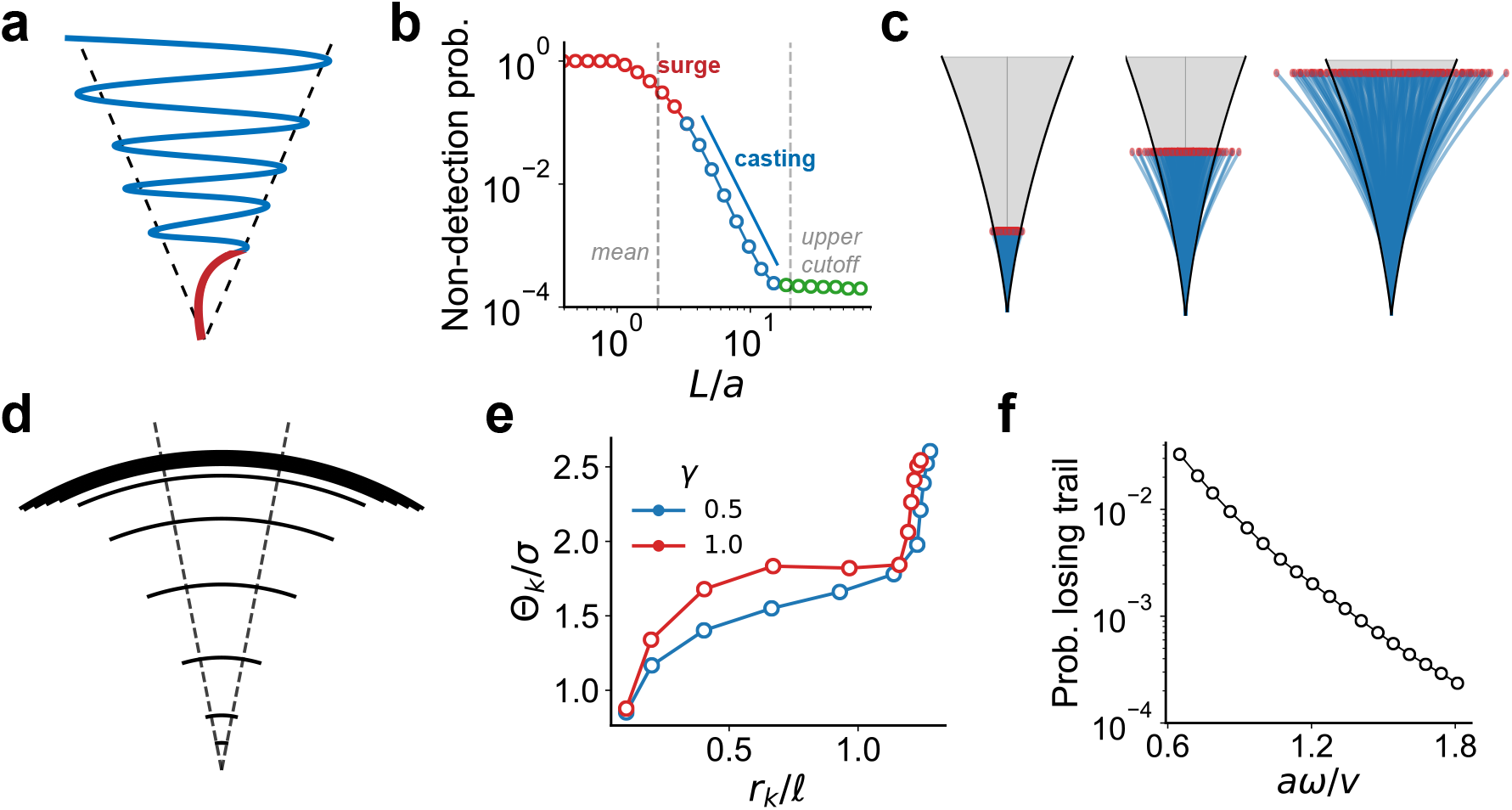
The role of casting in trail tracking. (a,b,c) The probability of not detecting the trail at distance *L* decays exponentially during the initial forward surge (red) and as a power-law during casting (blue) over a conical envelope. The searcher often finds the trail within the initial surge (mean in panel b) whereas casting determines the probability of losing the trail (determined by the upper cutoff) if the trail is not found during the surge. Beyond a characteristic lengthscale *ℓ*, the detection-rate becomes negligible (green region in panel b) as trails initially well-inside the casting envelope “escape" out of the envelope (panel c). (d,e) The casting policy obtained from Bellman optimization. The specific casting strategy depends on trail statistics. Note the increasing casting angle and the slowing down of the agent at *r* ≈ *ℓ* (panel e). (f) The trade-off between the probability of missing the trail for fixed 〈*L*〉 and searcher speed *v*.

A number of transformative experimental assays are suggested by our theoretical framework. The broad theme is whether and how animals adapt their behavior to the statistics of trails. For field experiments, it would be informative to measure the statistics of natural trails, analogous to the statistics of natural images that has brought insight into the adaptation of visual responses^19–22^. Specifically, one can measure the auto-correlation of local heading and curvature of natural trails, which would test the validity and fix the parameters of our WLC-type models. In laboratory settings, the statistics of trails can be controlled by varying persistence, curvature or using broken trails (Figure 4a). The general issue of adaptation is articulated in the following four specific questions that stem from our work.

**Figure 4.**
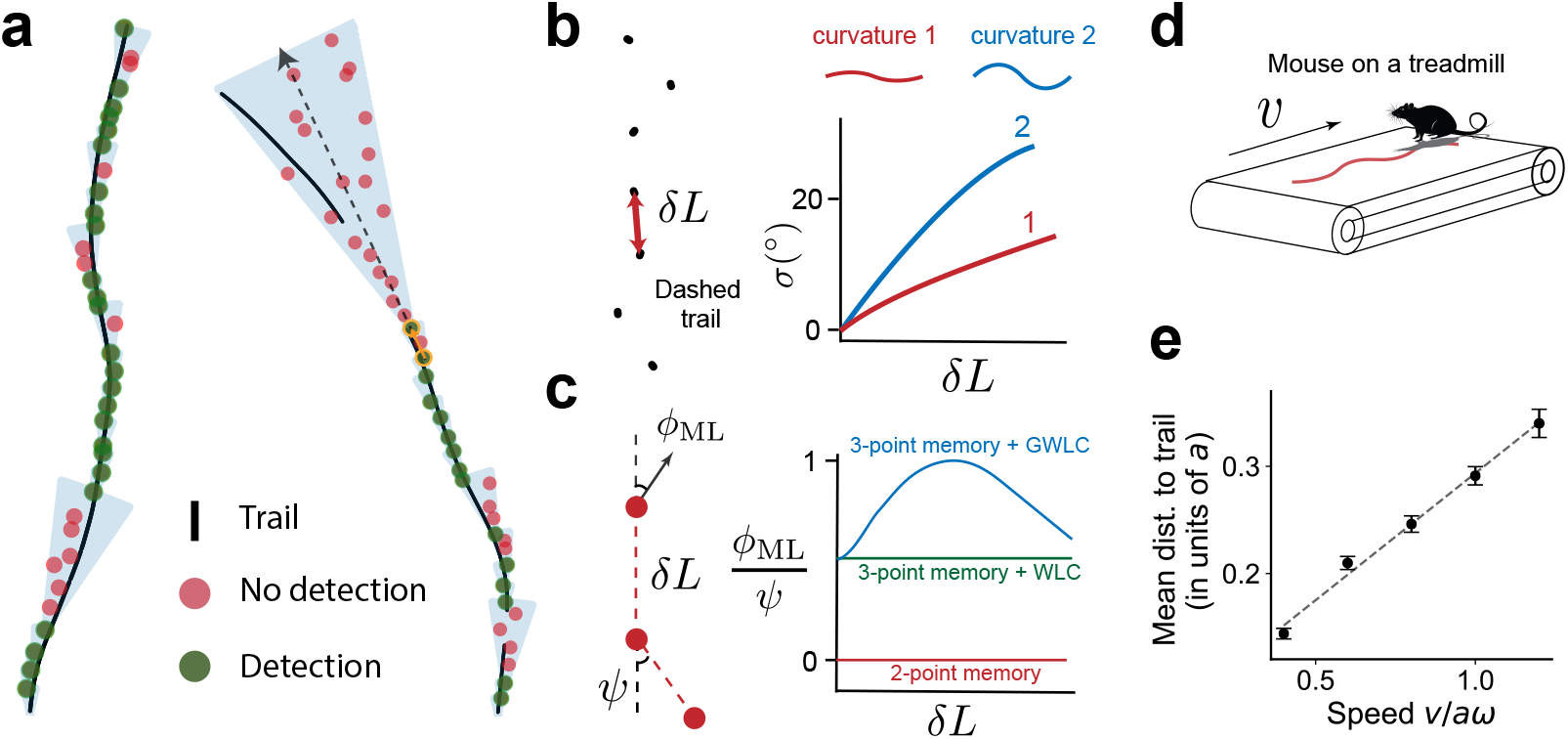
Behavioral assays to dissect trail tracking strategies. (a) A tracker executing a sector search often maintains continuous contact with the trail (left). Extended sector searches can instead be systematically elicited using broken trails (right). The subsequent search envelope’s orientation and width relative to past contact points informs the tracker’s internal estimate and uncertainty *σ* of the trail’s heading. (b) We propose dashed trails as an assay to infer how a tracking animal integrates past information. The distance between dashes forces contact points to be separated by at least *δL* and the subsequent search sector yields an estimate of *σ*. (c) Our theory predicts that the most likely trail heading *ϕ*_ML_ is proportional to the angle *ψ* between the line segments joining points of contact (red dots), with a pre-factor that depends on the distance *δL* between the two most recent contact points, the trail model and memory. (d) Automated behavioral tracking of rodents on a treadmill allows control of tracking speed and trail statistics^5, 18^. (e) The mean distance to trail with speed in simulations of sector search (as in panel a (left)), which recapitulates the linear relationship (dashed line) observed in experiments with rats^5^. We use a sector search strategy where the longitudinal speed v is fixed and the tracker rapidly casts within a conical envelope (see Figure M4a for a sample trajectory in a single bout). For each *v*, we simulate 100 trials, where each trial consists of successive 10 successful contacts with the trail. Trails are generated from the GWLC ensemble with *κ* = 0, *ξ*/*λ* = 3, *a*/*λ* = 0.1. Error bars are 1 s.e.m.

First: how long after the loss of contact do animals “give up” tracking? Our prediction is that they should when they get beyond the characteristic correlation length of the trails. At this point, the value of past information has expired and it is best to turn back or start a new search. This prediction can be tested by varying trail statistics, interrupting the trails and measuring when animals give up.

Second: does the amplitude of casting depend on the statistics of trails and the inter-detection interval? We predict that it should, and the specific quantitative relationship is a signature of the underlying predictive model employed by the animal (Figure 4b). Our prediction should be contrasted with the non-adaptive casting envelope assumed in ref. 5. The IDI can be experimentally manipulated by generating dashed trails as shown in Figure 4b, which forces the animal to detect the trail sporadically yet at controllable intervals. It would be particularly informative to verify whether or not animals include curvature in their estimates of the trails’ future heading or limit to persistence.

Third: what is the memory of past trail contacts? Experiments with forked trails^5^ show that rats exhibit a predictive component, suggesting a memory that extends over the recent past. Our theory posits that the tracker remembers (at least) the two most recent detection points. For the case of two-point memory, we predict the sector search is oriented along the line connecting those two points. If more than two points are remembered, the expected heading deviates from this line (by an angle that we calculate explicitly in the Supplement) as illustrated in Figure 4c. Note that the heading is not an average of the past headings, as assumed in ref. 5, and actually depends on the IDI between recent contacts: this prediction could again be tested by using curved, dashed trails as in Figure 4b.

Fourth: does the tracking speed vary with the typical IDI, reducing with increased uncertainty as predicted by the speed-accuracy trade-off Eq. (2)? This can be tested by varying the speed, for instance of the treadmill in ref. 5 (see Figure 4d), and measuring tracking accuracy. Available data for three speeds in ref. 5 are captured by our theory (see Figure 4e), which highlights the importance of revisiting those pioneering experiments and measuring additional quantities, namely the explicit prediction in Figure 3f.

In conclusion, we show that an optimized sector search strategy based on the memory of two or more recent detection events yields an oscillatory search path with increasing amplitude that naturally unifies the observed low-amplitude “zig-zagging” and larger-amplitude “casting” behaviors into the same quantitative framework. This framework elucidates the geometric and computational constraints faced by tracking animals, and identifies general features of the algorithms that efficiently solve the task, which can also be implemented for robotic applications. Insights and predictions developed here impact and should inform the design and analysis of future animal behavior experiments.

## METHODS

### A reinforcement learning framework for trail tracking

Trail tracking naturally splits into discrete episodes where, after each loss of contact, the tracker searches and attempts to re-establish contact with the trail. We use reinforcement learning (RL) to identify optimal search strategies for each episode and explore how factors such as sensory configuration or movement constraints influence the strategy. The task is a onedimensional search over angular space *θ*, the geometry of which is illustrated in Figure 2a. The tracker controls its tangential speed 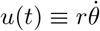 while its radial speed *v* is kept constant. For simplicity, we focus here on the configuration featuring a single sensor of size *a* sampling with a Poisson frequency *ω*. The generalization to two sensors is found in SI.

In each episode, the (Bayesian) agent maintains a posterior probability distribution function (PDF) P(*ϕ*) over possible trail headings, which is continuously updated based on the locations already visited, until the trail is recontacted. The agent’s strategy of decisions about its future trajectory *a priori* depends on the full high-dimensional distribution P(*ϕ*), which is difficult to learn. To circumvent this issue, we formulate the search task using a tractable parametrization of the posterior as a mixture of *K* Gaussian basis functions:

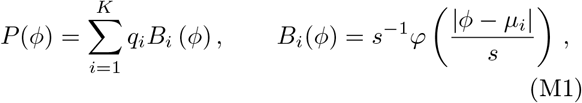

where φ is the standard normal PDF and the *q_i_*’s are normalized weights. The posterior is encoded by the weights ***q***, the posterior probabilities of the latent states given the agent’s history. The corresponding vector is lower-dimensional and yields to standard RL methods. For simulations in the main text, we used *K* = 3, equal initial weights, *μ*_1_, *μ*_2_, *μ*_3_ = −*σ*, 0, *σ*, and *s* = 0.5*σ*. These values were chosen so that the prior has mean zero and standard deviation ≈*σ*. Using *K* > 3 led to similar strategies of search but training was slower. We define the detection probability given latent state *i* for an agent at ***r*** as

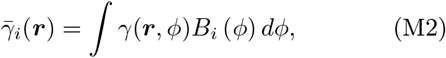

where, *γ*(***r***, *ϕ*) is the detection probability of finding a trail headed along *ϕ* if the searcher is at ***r***. We assume a Gaussian detection kernel of size *a*, with distance measured to the closest point on the trail: 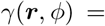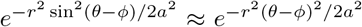, where we have used the small-angle approximation. Conditional on no detection at ***r***, Bayes’ rule yields 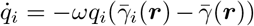 where 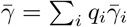 is the total probability of detection.

**Figure M1.**
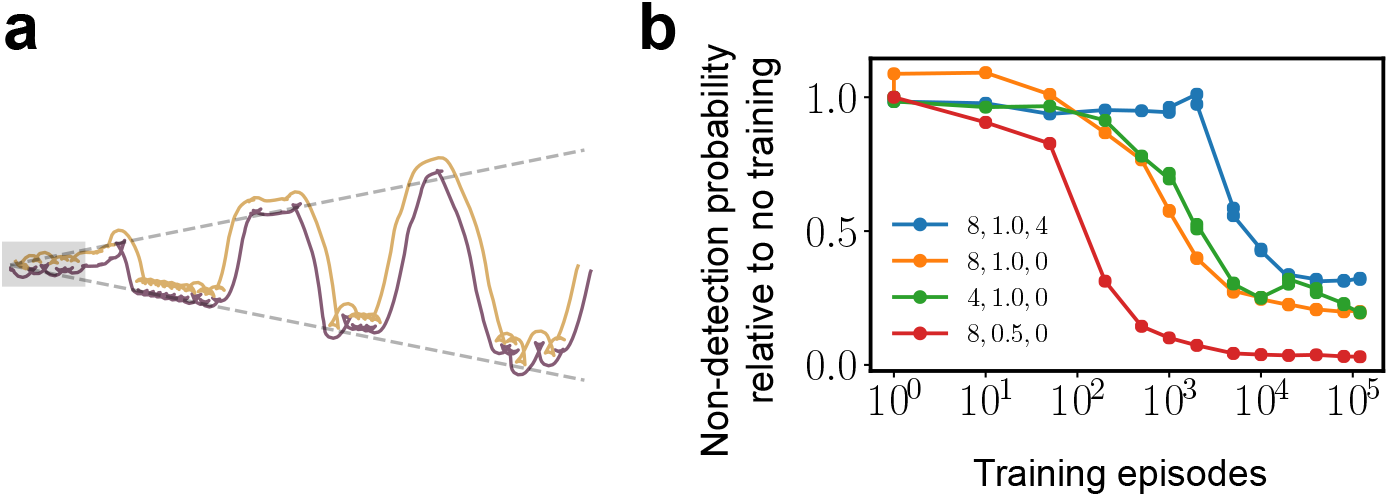
Two-sensor RL sector search path and learning curves. (a) The complete search path for an RL agent with two sensors. The gray shaded region close to the last detection point is shown in Figure 1f to highlight the criss-crossing pattern. At large distances, the search path shows casting similar to the one-sensor case (Figure 1e). (b) The probability of not finding the trail until time T (set to 100) as a function of the training time normalized by the probability of not finding the trail without any training. We show results for various values of the parameters (*α*, 1/*β*, *d*), where *α* ≡ *u*/*αω*, *β* = *aω*/2*vσΘ*_0_, and 2*d* is the distance between sensors for the two sensor case (see SI). The curves are meant to demonstrate non-trivial learning for different conditions.

From (M2), we have

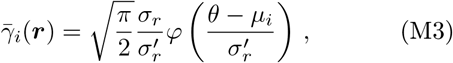

where *σ*_*r*_ = *a*/*r*, and
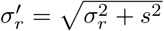

We use a discount rate *λ* and provide a reward as discussed below, after (M4). Training is performed in an episodic fashion with each episode lasting time *T*. The kinematic variables are updated with timestep dt and actions are taken with timestep *dt_act_* ≥ *dt*. Movement constraints are imposed by restricting the set of actions to three values, *u*/*aω* ∈ {-*α*, 0, *α*}. The state-space has four dimensions, *r*, *θ*, −ln *q*_1_/*q*_2_ and −ln *q*_3_/*q*_2_. We discretize our state space using a non-overlapping tile coding scheme^23^. We refer to the SI for hyperparameter values, details about the state space architecture and the case of two sensors.

We use a SARSA *Q*-learning algorithm^23^, which learns the so-called *Q* function, that is the value function for each action in a given state:

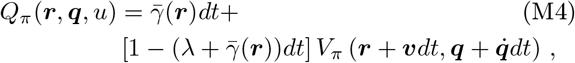

where *V*_*π*_(***r***, ***q***) = Σ_*u*_ *Q*_*π*_(***r***, ***q***, *u*)*π*(*u*| ***r***, ***q***), and the index *π* highlights the dependence on the probabilistic policy *π*(*u*| ***r***, ***q***). The above equation differs from the standard SARSA update by the addition of 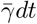 in the discount term, which is due to our formulation of the search as a continuing process conditional on no contact with the trail. Alternatively, one may provide a unit reward when the trail is found, stop the episode and start over. However, the credit assignment problem in goal-oriented tasks makes the training (see (M5) hereafter) problematic even though the final optimal policy is equivalent (see, e.g.,^24^). Our formulation circumvents both issues by (i) giving a local reward 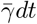 rather than a final one, which addresses the credit assignment, and (ii) including the detection probability 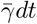 into the discount rate to account for the condition of no contact.

To learn the *Q*-function as defined in (M4), we use a “softmax” training policy, i.e., 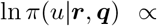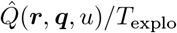 where 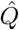 is the current estimate of the *Q* function, and *T_explo_* is a “temperature” parameter that is annealed as training progresses to allow for sufficient exploration of actions. Given an action *u* at state (***r***, ***q***) and a subsequent action *u′* at state (***r**′*, ***q**′*), 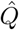 is updated during training as

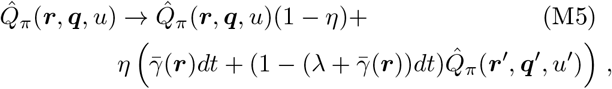

where *η* is the learning rate. The function 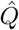 obtained at the end of the training period yields a search strategy as: 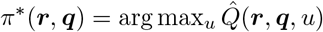.

We applied the algorithm just described for a range of values of *v*, *α* and *d* (half the distance between sensors for the two-sensor case). For each parameter set, we obtain a search strategy, the corresponding probability 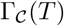 of missing the trail in time *T*, and the expected number of samples to find the trail. Comparing 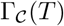 on a test set for different number of training episodes (Figure M1b) shows that non-trivial learning takes place, typically saturating at ~10^4^ training episodes.

### Casting in a minimal model of sector search

In order to establish the relation between casting and marginal value theory, here we propose a minimal model of sector search. The model lends to an analytical solution which allows us to quantify how the frequency of casting and the efficiency of search depend on the movement and computational constraints imposed upon the tracker.

We consider the same setting as the above episodic RL framework, where the tracker is searching for the trail over a sector after losing contact with it. To focus on casting, we analyze the behavior of the searcher after an initial forward excursion along the most likely heading. This surge decreases the probability weight *q* of the mode at *ϕ* = 0 and yields a symmetric bimodal posterior distribution concentrated at the two modes, ±*ϕ*_0_ (typically *ϕ*_0_ ~ *σ*). The resulting model is equivalent to (M1) with *K* = 2, μ_1_ = −*ϕ*_0_, μ_2_ = *ϕ*_0_ and *s* small. The searcher moves radially as *r*(*t*) (fixed), controls its tangential speed 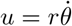, receives a unit reward when it finds the trail and incurs a movement cost per unit time, *μu*^2^/2, where *μ* sets the movement constraint. The reward and cost are discounted at a rate *λ*. The two-dimensional state space of the agent consists of *θ* and the probability *q* of finding the target at *ϕ*_0_ (the probability is 1 − *q* at −*ϕ*_0_). For full details, we refer to the SI.

The above model is exactly solvable. An optimal searcher exhibits oscillations between −*ϕ*_0_ and *ϕ*_0_ (“casting”) until it finds the trail (Figure M2a). After an initial transient, the searcher traverses a loop in state space (*θ*, *q*), alternating between sampling at −*ϕ*_0_ or *ϕ*_0_, and casting to the other side:

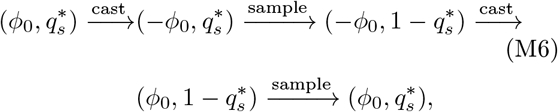

where 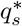 is the optimal switching probability at which the searcher stops sampling at *ϕ*_0_ and traverses to −*ϕ*_0_.

The speed of traversal from *ϕ*_0_ to −*ϕ*_0_ is determined by balancing the cost of traversing quickly and the potential value at −*ϕ*_0_ discounted due to the limited time horizon.

Intuitively, the searcher casts when the marginal value of continuing to sample at *ϕ*_0_ is *just* outweighed by the marginal value that the searcher receives if it stops sampling, traverses from *ϕ*_0_ to −*ϕ*_0_ and samples at −*ϕ*_0_. Balancing the marginal value of these two possibilities then yields the optimal switching probability 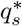. For small 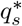, we derive (see SI):

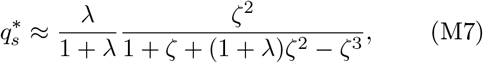

where 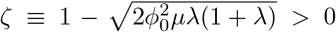. When ζ < 0, the movement cost outweighs the value the agent may receive, and the optimal strategy is to simply not move. 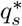 get smaller and thus the searcher casts less frequently with increasing time horizon (*λ* ≪ 1) or movement costs (0 < ζ ≪ 1). The probability of not detecting the trail after time *t* is given by

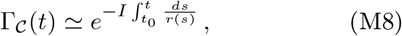

where *I* is interpreted as the rate of information acquisition. Its dependence on μ, *λ* and *ϕ*_0_ is shown in Figure M2b. As expected, increasing movement cost (decreasing ζ) decreases how quickly information is acquired. Similarly, a large time horizon (*λ* small) makes the agent sample at ±*ϕ*_0_ longer and cast slower, decreasing the rate of information acquisition. For a constant radial speed *v*, the probability 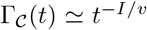 decreases as a power-law, which arises quite generally from the sector search geometry as discussed in the main text.

### Trail statistics

If an agent makes contact with the trail at two points separated by distance L, statistical and geometric information is encoded in the “propagator” *P*(*ϕ*_L_, *ϕ*_0_), where *ϕ*_L_, *ϕ*_0_ are the trail headings at the two points (measured relative to the line joining them). If *H* points of contact are remembered, then the propagator can be used to compute the posterior distribution of the heading: 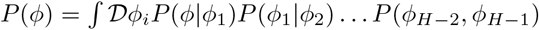. We introduce an ensemble of trails (which we call the Generalized Worm-Like Chain (GWLC) ensemble) that have persistence in heading and curvature, quantified respectively by the parameters *κ* and *λ* (distinct from the discount rate used for RL) and a typical radius of curvature, *ξ*. Typical samples of trails from this ensemble are presented in Figure M3a. The WLC ensemble^16, 17^ previously introduced for polymers is a special case with *ξ* = ∞. The propagator *P*(*ϕ*_L_, *ϕ*_0_) takes into account all the trails constrained to pass through two contact points, weighted by their probability. In particular, we define

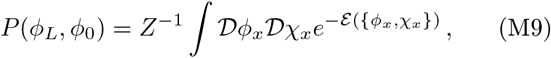

where *Z* is a normalization constant and the integral is over all possible headings and curvatures *ϕ*_*x*_, *χ_x_* at the various positions *x*. The trails satisfy the constraints *y*_0_ = *y_L_* = 0 and have end-point headings *ϕ*_0_, *ϕ*_L_. The action 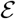 of a path is given by

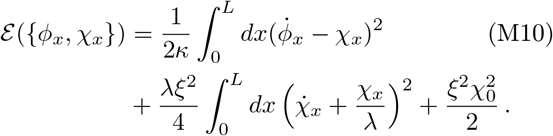

**Figure M2.**
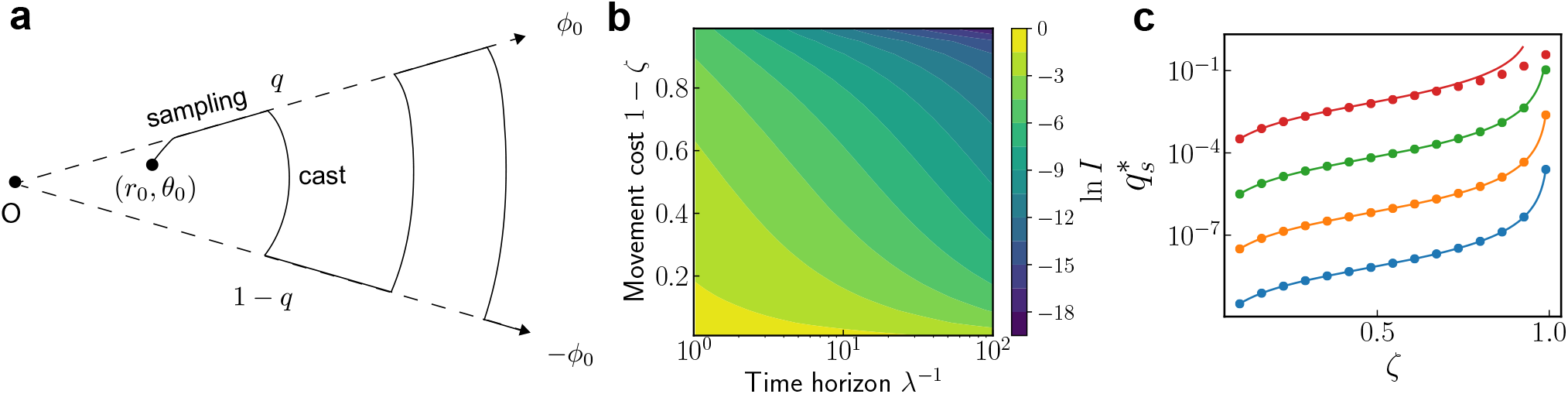
(a) The behavior of the agent searching for the trail which is either along −*ϕ*_0_ or *ϕ*_0_. The agent alternates between sampling at ±*ϕ*_0_ and traversing to the other side. *q* is the agent’s estimate of the probability that the trail is along *ϕ*_0_. (b) The rate of information acquisition *I* as a function of the movement cost (1 - *ζ*) and the effective time horizon (*λ*^−1^). (c) The optimal switching probability, 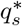, as a function of *ζ* for *λ* = 10^−8^, 10^−6^, 10^−4^, 10^−2^ plotted with blue, orange, green and red respectively. The dots are the exact solution obtained numerically (see SI), and the solid lines are from using the approximation (M7).

Here, we have oriented the x axis along the line joining the two points of contact and used the small-angle approximation so that 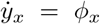. The model (M10) is a Gaussian process. Since symmetry dictates that 〈*ϕ*_L_〉 = 〈*ϕ*_0_〉 = 0, it follows that *P*(*ϕ*_L_, *ϕ*_0_) is defined entirely by the variance 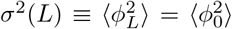, and by the correlation 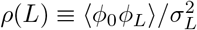.

We present the full calculation of *σ*(*L*) and ρ(*L*) in the SI. In summary, the first integral over χ_*x*_ can be performed using the Gaussian integral formula and leads to an effective action in *ϕ*_*x*_. The Euler-Lagrange equation of this effective action then yields extremal paths (“splines”) that minimize the effective action (see SI for details). The splines have the form:

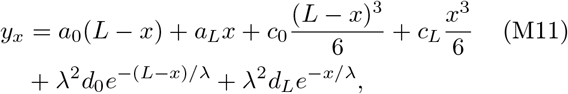

where the constants in the above equation are set so that *y*_0_ = *y_L_* = 0 and can be expressed in terms of *ϕ*_0_, *ϕ*_*L*_. Figure M3b shows the splines between contact points spaced at increasing intervals. Plugging (M11) in the effective action, we obtain

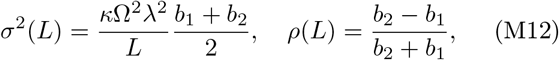

where the two functions *b*_1_ and *b*_2_ are

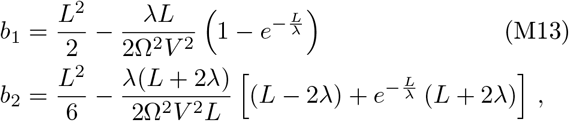

and 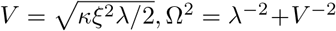. The variance *σ*(*L*) and the correlation *ρ*(*L*) are plotted in Figure M3c,d. Three distinct regimes are apparent. For *L*/*λ* ≪ 1, we can appproximate *σ*^2^(*L*) ≈ *κL*/3 + *L*^2^/4*ξ*^2^, which reflects diffusive ∝*L* and curvature-dominated ∝*L*^2^ scalings. When diffusion dominates, *ρ*(*L*) = −1/2, whereas *ρ*(*L*) ≈ −1 when curvature dominates. The perfect anti-correlation in the curvature-dominated regime is intuitive as the line joining the two points of contact can be viewed as the chord of a circle with radius *ξ*, whereas *σ*(*L*) = *L*/2*ξ* is the angular deviation of the trail around this chord. When *L*/*λ* ≫ 1, the heading is randomized over many correlation lengths and the diffusive scaling *σ*^2^(*L*) ≤ 2*λL*/3*ξ*^2^ is recovered.

While *P*(*ϕ*_*L*_|*ϕ*_0_) (the interpolation model) is required to integrate past information, search strategies also require the *forward* propagator *P*(*ϕ*_*L*_, *y_L_* | *ϕ*_0_), which keeps track of trail headings and locations while the agent searches for the trail. The methods described above can be used again (see SI for details) to yield extremal paths as (M11), and the three quantities 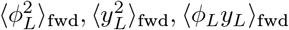. These quantities fully describe *P*(*ϕ*_*L*_, *y_L_* |*ϕ*_0_). We validated our interpolation model (M12) and the forward model using numerical simulations (Figure M3e,f).

**Figure M3.**
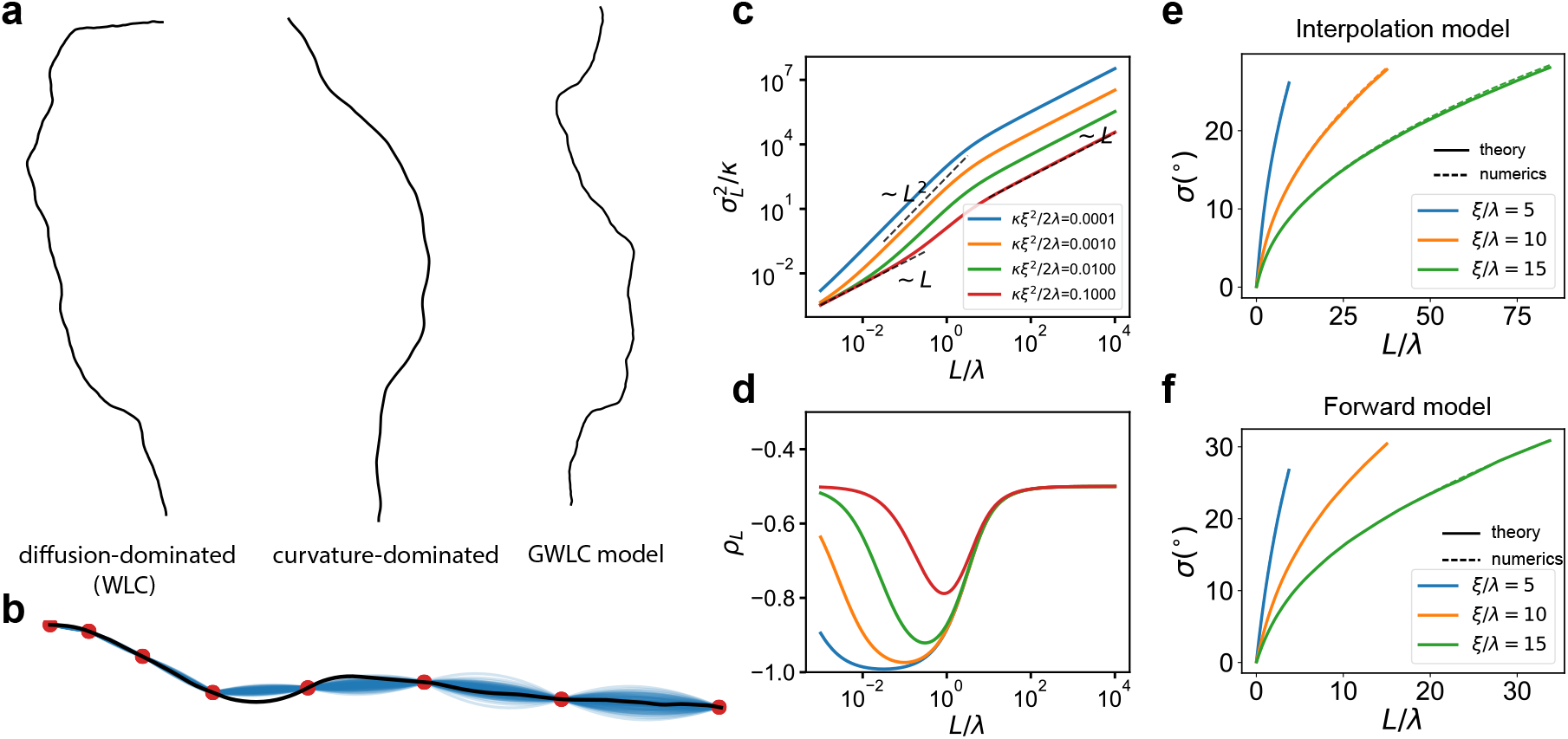
(a) From left to right: samples of trails diffusion-dominated (*κλ* = 0.05, *ξ*/*λ* = ∞), curvature-dominated (*κλ* = 0, *ξ*/*λ* = 7), and trails which have both diffusive and curvature components (*κλ* = 0.05, *ξ*/*λ* = 7). (b) Extremal paths (blue) between detection points (red) on a trail (black) are plotted using (M11) for *kλ* = 0, ξ/*λ* = 7. (c,d) The uncertainty 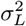 (plotted in log-log scale to highlight the three scaling regimes) and the correlation *ρL* between end-point angles versus the distance between detections, *L*, for the generalized worm-like chain (GWLC) model. (e,f) Numerical validation (dashed lines) of the theoretical predictions (solid lines) obtained by generating trail samples from the GWLC ensemble.

### The non-detection probability during surge and cast

We introduce a sector search strategy that allows us to quantify the non-detection probability taking into account the full dynamics of the trails and yields intuition on the factors that contribute towards losing the trail entirely. We suppose the radial speed *v* is fixed and the tangential speed *u* ≫ *aω*. In other words, the agent casts rapidly within a conical envelope of semi-aperture angle *σ*Θ_0_, where *σ* is the prior uncertainty of the trail’s heading and Θ_0_ ≳ 1 (Figure M4a). To simplify the presentation, we assume curvature-dominated trails, i.e., *σ*(*L*) = *L*/2ξ, though the arguments below are general. The rapid casting limit *u*/*aω* ≫ 1 allows us to compute the non-detection probability at distance *r* from the most-recent contact point defined by Eq. (3):

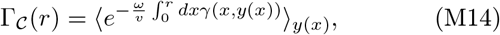

where the expectation is over the full ensemble of trails {*y*(*x*)} that pass through past detection points. Here, we provide intuition for the detection rate *ωγ*(*x*, *y*(*x*)) and refer to the SI for full details. For small *x*, i.e., *x* ≲ *a*/2*σ*Θ_0_, *γ*(*x*, *y*(*x*)) = 1 for most trails as the sensor size a spans the entire casting envelope 2*σx*Θ_0_. In the casting regime (*x* ≳ *a*/2*σ*Θ_0_), since the time spent on the trail in a single cast is *a*/*u*, the probability of non-detection per crossing is *e*^−*aω*/*u*^. After n crossings, the non-detection probability is then e^−*naω*/*u*^. As the agent moves a distance *dx* in the radial direction, it crosses the trail *n* = *udx*/2*vσx*Θ_0_ times, i.e., *γ*(*x, y*(*x*)) = *a*/2*σx*Θ_0_, which is independent of *u*. From (M14), these relations yield an exponential 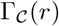 during the initial surge followed by a power-law, *r*^−β^ with exponent *β* ≡ *aω*/2*σv*Θ_0_. This heuristic argument aligns well with the results of numerical simulations in Figure M4b.

The spline formulation of the GWLC yields a geometric picture of the dynamics. Intuitively, the nondetection probability (and thus the posterior probability from Eq. (1)) is large if the trail has a large probability of escaping the casting envelope before it is found by the tracker. In order to “escape” the casting envelope, the trails that are initially well within the casting envelope have to bend significantly. This bending incurs a cost in the action (M10), which reduces with increasing *r*. The non-detection probability conditional on initial trail heading, 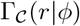, flattens out (i.e., the detection rate vanishes) when a significant fraction of trails escape the casting envelope, including trails with initial heading *ϕ* = 0 (Figure 3c).

**Figure M4.**
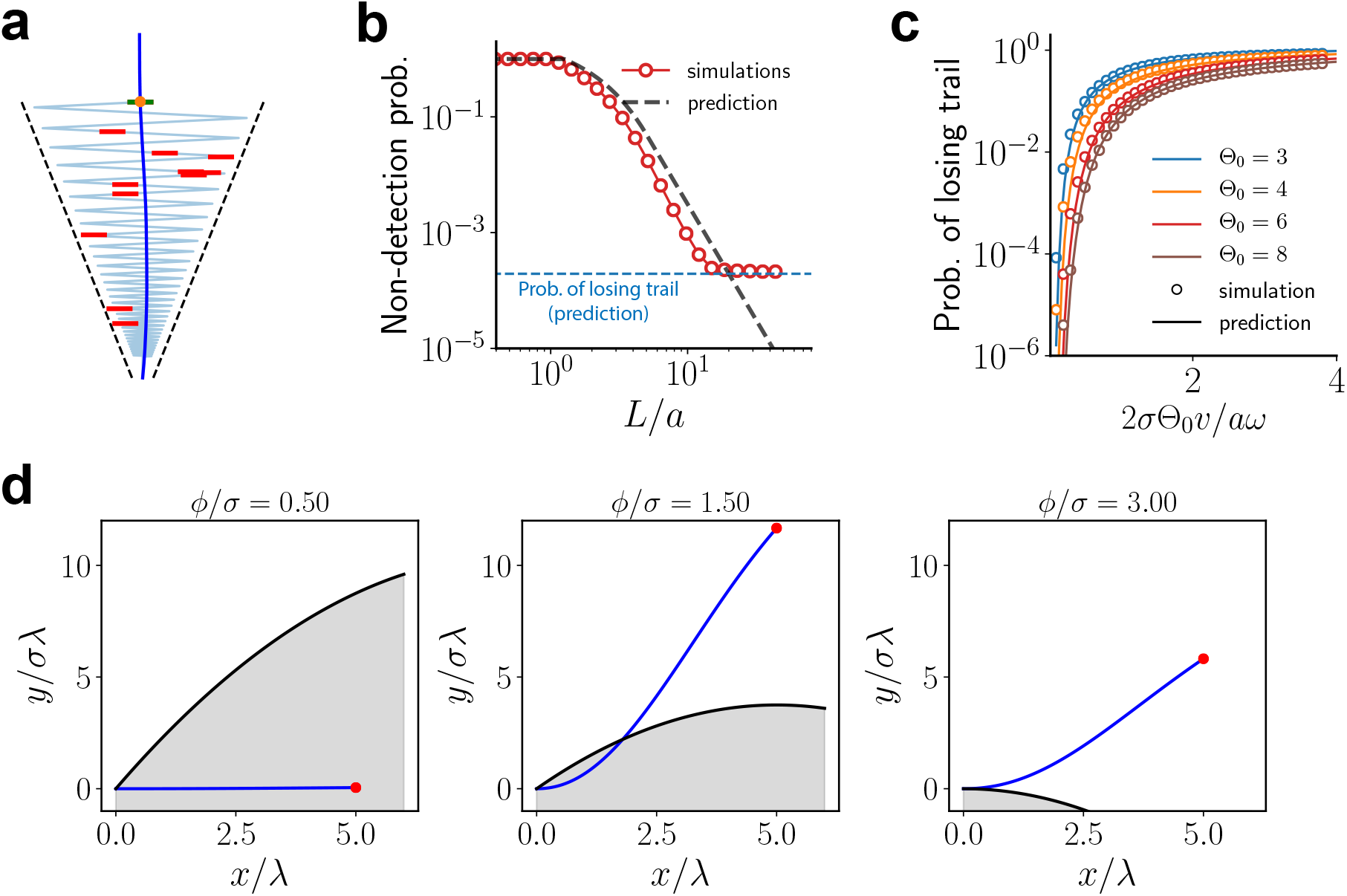
Sector search over a conical envelope: simulations. (a) A sample episode of a sector search starting from the most recent point of contact with the trail in dark blue. The tracker samples at discrete locations (red) with Poisson statistics while casting within the conical envelope until it re-establishes contact. (b) The prediction (black dashed line) of an exponential non-detection probability during the surge followed by a power-law decay during casting aligns well with simulations in panel (a) (red circles). Parameters are 1/*β* ≡ 2*σv*Θ_0_/*aω* = 0.25, Θ_0_ = 5, *σ* = 0.02. Trail parameters are *κ* = 0, *ξ*/*λ* = 5, *a*/*λ* = 0.1 for all panels. (c) The probability of losing the trail in simulations (open circles). The solid lines correspond to the theory prediction 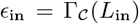, where *L*_in_ = 2*σξ*Θ_0_ (see Methods). (d) The path of the trail (blue) that is most likely to go undetected for three values of *ϕ*. The black line is the rotated casting envelope *R*(*x*) = *σx*Θ_env(x)_ − *xϕ*, *σ* = 0.17(10°).

This geometric picture yields a length scale *L*_in_, which is the distance at which the trails initially along the most-likely heading *ϕ* = 0 escape the casting envelope. For curvature-dominated trails, escaping trails should deviate by an angle *σ*Θ_0_ ≃ *L*_in_/2*ξ*, which gives *L*_in_ ≃ 2*σξ*Θ_0_. The probability of losing the trail can then be estimated as 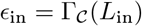. An additional contribution to the flattening of 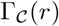 comes from the trails which are headed outside of the cone |*ϕ*| > Θ_0_ even before reaching *L*_in_. The probability 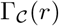 would then flatten out at a different length scale L_out_, where 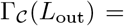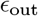. The relative contributions of the two mechanisms are discussed in detail in the SI and they are validated by using numerical simulations as shown in Figure M4c and Figure M5b.

Due to the power-law tail for 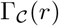 with exponent *β*, the mean distance to find the trail, 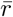, is determined by the upper or lower cutoff for *β* < 1 or *β* > 1, respectively. Since the upper cutoff is much larger than the lower cutoff, it is more convenient for the tracker to use *β* > 1. By using the self-consistency condition 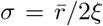 in the curvature-dominated regime, and the relation 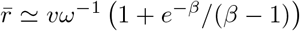, we obtain for the maximum speed

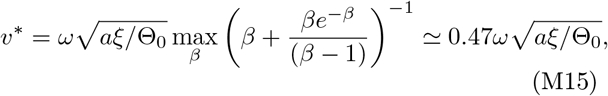

where the maximum occurs at *β*\* = 1.63 (see Figure M5c). The mean number of samples to find the trail at this optimum is ≈1.3, indicating that the optimal strategy is for the agent to move slow enough that it typically finds the trail within one or two samples.

Intuition for the above results holds quite generally, including for other statistical ensembles of trails and for non-conical casting envelopes. The latter is relevant because we expect that a slow down in the radial direction and a widening casting envelope may increase the likelihood of finding the trail. Relaxing the constraints of fixed radial speed and conical envelopes constitutes the aim of the next Section.

**Figure M5.**
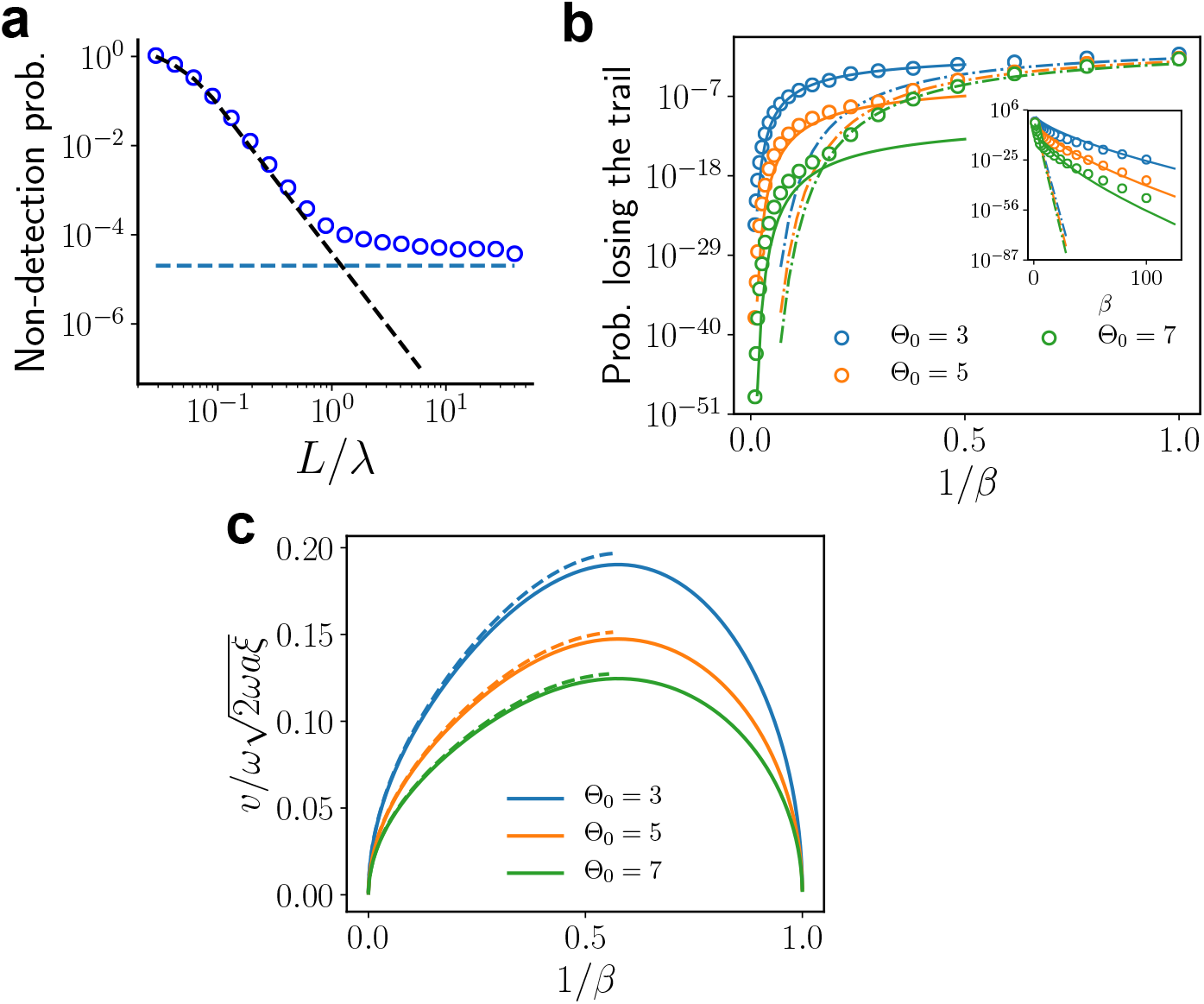
Sector search over a conical envelope: numerics. (a) The non-detection probability obtained by numerically evaluating (M14) (blue circles) instead of simulating the process (Figure M4a). The black and blue dashed lines are the theory predictions as in Figure M4b. (b) The probability of losing the trail computed numerically (circles). The dash-dot lines correspond to predictions (to a constant prefactor of order one) from the regime where the missed trails escape the cone, *ϵ*_in_, and the solid lines corresponds to the prediction when the trails outside the cone are the ones most likely to be missed, *ϵ*_out_. Inset shows the same data plotted against *β* ≡ *aω*/2*σv*Θ_0_ to highlight the small *v* behavior. (c) The speed obtained from imposing the self-consistency relationship between IDIs and uncertainty (see Eq. (2)). Solid lines are from theory in the curvature-dominated limit of the GWLC ensemble and dashed lines are from the values that satisfy the self-consistency relationship computed numerically.

### Bellman optimization of the casting strategy

Our final step is to formulate the problem of optimizing the casting strategy. The geometry of the search is shown in Figure M6a. To simplify, we parametrize the path by the set of turning points {*r_i_*, *θ*_i_}. We assume a fixed speed v (note that the average *radial* speed depends on the strategy), which is set later based on the probability of losing the trail. As discussed in the main text, we optimize the average tracking speed, 〈*L*/*T*〉, after constraining the inter-detection interval, 〈*L*〉, using a Lagrange multiplier Λ (see Eq. (4)). This optimization can be recast as a Bellman-type dynamic programming problem by breaking the problem up into discrete steps, each corresponding to a cast from {*r_i_*, *θ*_i_} → {*r*_*i*+1_, *θ*_i+1_} (denoted {*r′*, *θ′*} → {*θ*} below for conciseness):

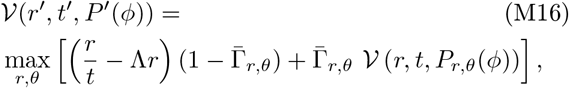

where the required reward function in Eq.(4) is 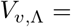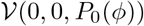 with the Gaussian normal prior 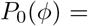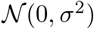. The first term in the square bracket returns the reward if the trail is found (weighted by the detection probability 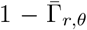, while the second term advances to the next cast if detection failed. The Bellman equation thus relates 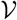 before and after a cast by updating the current state variables, *r′*, *t′*, *P′*(*ϕ*). The time elapsed is updated as *t* → *t′* + (*r*, *r′* + 2*r*|*θ*|)/*v*, where we approximate *θ*^0^ *θ* 2 *θ* for simplicity. The prior *P*^0^(*ϕ*) is updated using Bayes’ rule 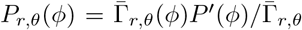, where 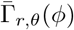 is the non-detection probability given the initial trail heading *ϕ*, and 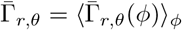 is the normalization.

**Figure M6.**
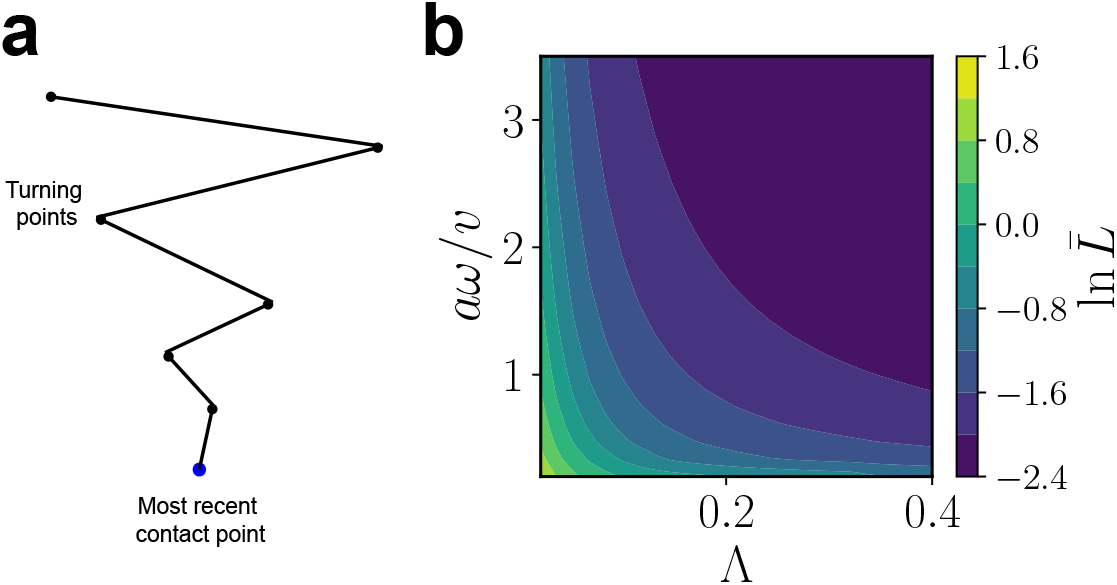
We approximate a zigzagging casting trajectory as a sequence of casts parameterized by turning points {*r*_*i*_, *θ*_*i*_}, which are defined using polar coordinates with the origin at the last contact point. The turning points are optimized using Bellman optimization so as to maximize the average radial speed 〈*L/T*〉 after constraining the average radial distance 〈*L*〉. (b) Contours of constant 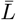 after optimization.

It remains to calculate 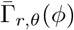, which depends on the *forward* model of the trail. We consider generally that the trail’s azimuthal position expands as *σ*_fwd_(*r*) i.e., a trail initially headed along *ϕ* is located at the azimuthal position, *ϕ* + Δ*ϕ*, where 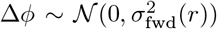. For the GWLC model, we show in the SI that *σ*_fwd_(*r*) = *σ*(*r*), where *σ*(*r*) is given in (M12). Since the time spent on the trail within the casting envelope in a single cast is *a*/*v*, we have 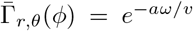 if |*ϕ* + Δ*ϕ*| < *θ* and equal to unity otherwise. Since Δ*ϕ* is normal-distributed, we have 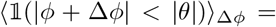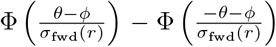, where Φ is the normal cumulative distribution function. We finally approximate 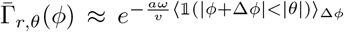, and compute the expectation over *ϕ* in 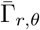. numerically. The previous approximation greatly simplifies the optimization as *P*(*ϕ*) is then a sufficient statistic for past measurements (thus allowing the decomposition in Eq. (M16)) and yet captures the effects of the trail’s widening trajectory on the optimized casting strategy.

At each casting step, we optimize (using standard black-box optimization methods in SciPy) for Δ*r* = *r* - *r*′ > 0 and *θ* by expanding Eq. (M16) two steps further into the future. The optimization then involves six variables, the immediate pair Δ*r*, *θ* and the subsequent two pairs. The optimized immediate pair is then used for updating as detailed above, and the process is repeated. Optimizing more than two steps did not yield different results as the 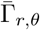 factors introduce an effective planning horizon, effectively suppressing contributions from future rewards beyond two steps. For less than two steps, optimization is “greedy” and the corresponding landscape was found to have a qualitatively different, shallow landscape around the optimum. After completing the optimization procedure, the constraint of fixed 〈*L*〉 is imposed by varying *v* and the Lagrange multiplier Λ (see Figure M6b).

## Acknowledgements

We thank Venkatesh Murthy for stimulating discussions throughout our work. This research was supported in part by NSF Grant No. PHY-1748958, NIH Grant No. R25GM067110, and the Gordon and Betty Moore Foundation Grant No. 2919.02. GR was partially supported by the NSF-Simons Center for Mathematical & Statistical Analysis of Biology at Harvard (award number #1764269) and the Harvard Quantitative Biology Initiative.

## Supplementary Information for ‘Sector search strategies for odor trail tracking’

## 1 Additional details of reinforcement learning implementation

Here, we expand on the details of the reinforcement learning implementation presented in the main text and Methods, specifically the case of two sensors and the hyperparameters used for training. We set *a* = 1, *ω* = 1 to fix the units.

The agent’s position vector ***r*** is updated with a time step *dt* = 0.05 and the agent’s actions are updated with a time step *dt*_act_ = 0.1. The agent chooses its tangential speed from three values, *u* ∈ 2 {-*α*, 0, *α*}, where *α* is a parameter. The tangential speed, *v*_tang_, is smoothed over a time scale *t*_smooth_:

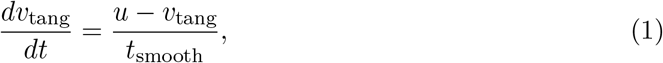

where we use *t*_smooth_ = 1. The agent’s heading is given by *θ*_agent_ = tan^−1^(*v*_tang_/*v*_rad_), where the radial speed *v*_rad_ = *v* is kept fixed. The two sensors are located at ***r***_1_, ***r***_2_ which are at a distance *d* from ***r*** in a direction perpendicular to the agent’s heading. The case of two sensors is similar to the case of one sensor described in the Methods except for the detection probability given latent state *i*. For one sensor, we have

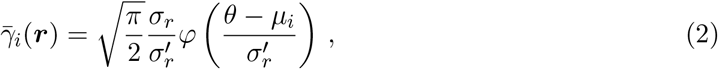

where *σ*_r_ = 1/*r*, and 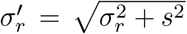. For two sensors, we assume that each sensor has a Gaussian detection kernel of size a and samples at a frequency *ω*/2, where the factor of half ensures that the two sensor case reduces to the one sensor case for *d* = 0. The detection probability is then 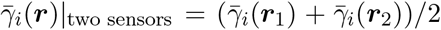. The rest of the details of the RL implementation remain the same as the one sensor case.

For the figures in the paper, we fix *σ* = 0.2 as the strategy depends largely on *σv*/*u*, which is varied by changing *σv* for a given *α*. The state-space has four dimensions, *r*, *θ*, – ln *q*_1_/*q*_2_ and – ln *q*_3_/*q*_2_. We discretize our state space using a non-overlapping tile coding scheme: the variable *r* is discretized using *σvT*/5 (rounded) states between 0 to *σvT*, *θ* into 15 states between −1.5*σ* and 1.5*σ*, and, finally, – ln *q*_1_/*q*_2_ and – ln *q*_3_/*q*_2_ into 5 states each between −4 and 4. We choose a number of samples *ωT* = 100, which implies that if *v*/*aω* = 1, for instance, we have 20 states for *r* and a total of 20 × 15 × 5 × 5 = 7500 states that tile the state space.

As noted in the Methods, we use a softmax policy with an annealing scheme for training. For each state *s*, we first normalize the *Q*-values: 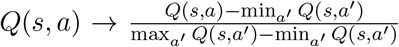 so that the normalized *Q* values lie between 0 and 1. The “temperature” parameter, *T*_explo_ is first set to 3 and annealed linearly down to 0.2 from the first training episode to the last training episode. For generating trajectories after learning, we set *T*_explo_ = 0.2. We use a learning rate *η* = 0.02 and discount rate *λ* such that 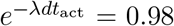 which corresponds to a time horizon *λ*^−1^ ≈ 5 samples.

## 2 Optimal stochastic control and searching

We first describe a general formulation of searching in the framework of optimal control and then proceed to analyze the case of sector search in more detail further below.

### 2.1 The Hamilton-Jacobi-Bellman equation for search problems

Consider an optimal control problem where the agent searches for a target at an unknown location ***y***. The agent has prior knowledge encoded in the vector ***ψ*** which is updated during the search. The agent receives a reward of one and stops searching upon finding the target. The probability per unit time of finding the target for an agent at ***x*** is given by *ωγ*(***x***, ***y***) with *γ*, being the detection kernel of size *a*. Additionally, there is a cost per unit time of controlling its velocity, 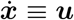, along various directions, μ*u*^2^/2 (*u* is speed and μ is some constant), and the rewards and costs are discounted per unit time using a discount rate *λ*. In this section, we set *ω* = 1, *a* = 1.

The agent has an internal model of how *ψ* is updated when it doesn’t detect the target at ***x***. In general, *ψ* represents all the information that the agent maintains about the location of the target. In particular, if we take as a posterior distribution, *P*(***y***), updated using Bayes’ rule starting from some prior, we have

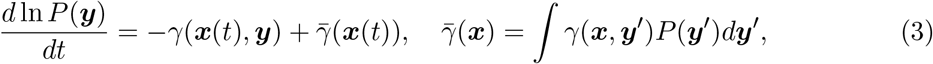

where ***x***(t) is the position of the agent at time *t*. Note that 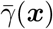 is the probability per unit time of detecting the target given current information. We introduce the value function *V* (***x***, *ψ*), which is the optimal expected sum of discounted rewards given the current position of the agent and available information about the target. The evolution of the value function in a time interval dt satisfies the dynamic programming equation

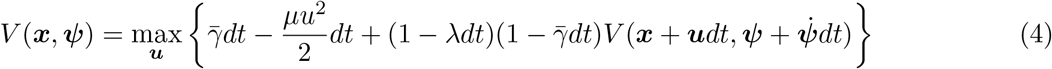

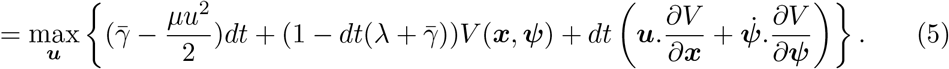

The first two terms on the right hand side of (4) are the probability of detecting the target and the cost of control in the interval *dt* respectively. The latter term is the discounted value function if the process continues, i.e., target is not detected in *dt*. The optimal policy is obtained by optimizing the terms in the curly brackets with respect to the control variables ***u***. Taking the gradient w.r.t ***u*** and equating to zero, we get

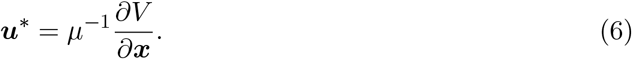

Plugging this back into (5) and canceling *V* (***x***, ***ψ***) on both sides, we get the Hamilton-Jacobi-Bellman (HJB) equation

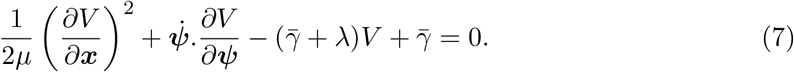

The solution of the above HJB equation yields the optimal policy via (6). In most cases, is high-dimensional and an exact solution is infeasible. We consider a minimal model of sector search which enables an exact solution, as described below.

### 2.2 A minimal model of casting

We consider a minimal model of trail tracking discussed in the Methods where the search is over a one-dimensional angular space. The model naturally gives rise to casting dynamics and quantifies how the frequency of casts and the efficiency of search depends on various movement and computational constraints.

Consider two point targets located at ±*ϕ*_0_. As noted in the Methods, these correspond to the modes of a bimodal posterior distribution of the heading of the trail after an initial forward excursion. Suppose q is the probability of the target at *ϕ*_0_. When the agent is at *θ* = ±*ϕ*_0_, we assume

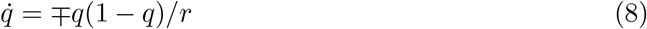

respectively, where the factor 1/*r* is the angle subtended by the detector at distance *r* from the last contact (note that we have set *a* = 1, *ω* = 1). For intermediate values of *θ*, 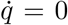. The radial path *r*(*t*) is assumed fixed a *priori* with *r*(0) = *r*_0_ > 0.

We would like to find the optimal control variable 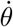 that maximizes the reward while minimizing a energy cost per unit time 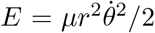. However, the dependence on *r* makes the problem non-stationary and challenging to analyze. To circumvent this issue, we introduce the coordinate 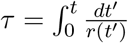. We have 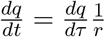 and 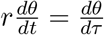, so that the update rule on *q* and the energy cost are both independent of *r* in the new coordinates. We use 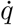 and 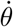 to denote the derivative w.r.t *τ* henceforth.

We re-write equation (4) for a one-dimensional search (along *θ*) with the above assumptions:

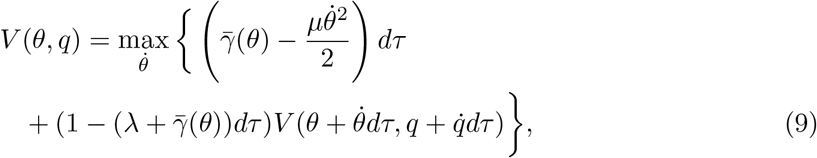

where, 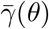 is the detection rate and *λ* is the effective discount rate in the new coordinates; if the time horizon in the t-scale is *T*, we have an effective time horizon 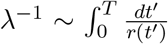 in the τ-scale. Taylor expanding the *V* in the curly brackets above to first order and optimizing for
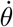 we have

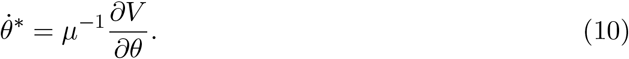

Plugging this back into (9) and canceling *V* (*θ*, *q*) on both sides, we get the HJB equation

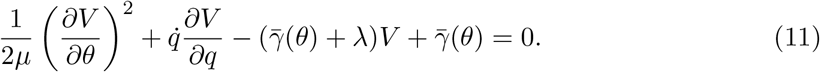

### 2.3 Solution to the HJB equation

The dynamics of the agent is as follows: 1) If the agent is initially between −*ϕ*_0_ and *ϕ*_0_, it chooses to travel to either *ϕ*_0_ or *ϕ*_0_ depending on its current *θ* and *q*. In other words, there is a location 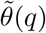, such that the agent travels to the right if the current position 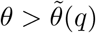 and vice-versa. 2) If the agent chooses to travel to *ϕ*_0_, say, then it travels at a speed that balances out the cost of traveling with the discounted value it expects to obtain at *ϕ*_0_. 3) Once at *ϕ*_0_, the agent begins sampling and if it doesn’t find the target, it remains stationary until *q* decreases to a ‘switching’ probability *q_s_*. At that point, a decision is made to traverse to −*ϕ*_0_. The choice of *q_s_* depends on the balance between the cost of traversing (at an optimized speed) and the potential future reward at −*ϕ*_0_ versus the expected reward from remaining at *ϕ*_0_. Once at −*ϕ*_0_, by symmetry, the agent samples until q increases to 1 − *q_s_* or until it finds the target. Eventually, the agent cycles between −*ϕ*_0_ and *ϕ*_0_ at an optimized speed, sampling for a fixed amount of time at each end, until it finds the target.

There are three choices made by the agent corresponding to the above three steps of the dynamics: 1) Given the initial *θ* and *q*, which direction should the agent move i.e., what is 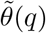? 2) At what speed should the agent move? 3) When should the agent switch from searching at one end to the other i.e., what is the optimal value of *q_s_*, 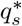? Below, we calculate 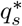 and the optimal speed. We do not calculate 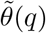, since it does not affect the agent’s long-term behavior of cycling between −*ϕ*_0_ and *ϕ*_0_.

We have 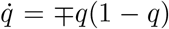 depending on whether *θ* = ±*ϕ*_0_, and 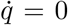 otherwise. Likewise, the detection rate is 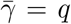, 1 − *q* at *ϕ*_0_, −*ϕ*_0_ respectively, and 0 otherwise. Since the agent remains stationary while sampling at *θ* = *ϕ*_0_, −*ϕ*_0_, we have ∂*V*/∂*θ* = 0 from (10) at these two points. Putting these together, (11) can be re-written as the set of equations

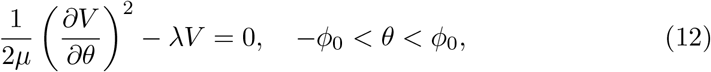

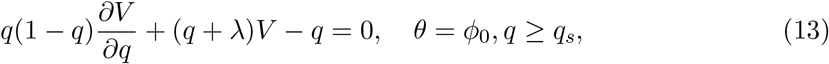

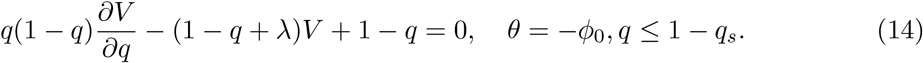

The first equation is the HJB equation of a “free particle” (with an additional discount factor) which describes the motion of the agent while it traverses from one end to the other. The latter two equations capture the evolution of the value function as the agent samples at *ϕ*_0_ and −*ϕ*_0_ respectively conditional on not finding the target.

The agent traverses a loop in state space:

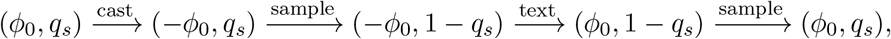

The solution is found by following the value function around this loop, obtained by integrating the above set of equations and matching the value function at (*ϕ*_0_, *q_s_*) at the beginning and the end of the loop. We exploit the symmetry *V* (*ϕ*_0_, *q_s_*) = *V* (−*ϕ*_0_, 1 − *q_s_*), which reduces the problem to matching the first and third points in the above loop. For an arbitrary choice of *q_s_*, the solution to the first equation yields the value function during a traversal between −*ϕ*_0_ and *ϕ*_0_, which is then used to match the value functions at the two boundaries obtained from the latter two equations. Optimizing over *q_s_* then yields the optimal policy.

−*ϕ*_0_ < *θ* < *ϕ*_0_: Consider the first equation for −*ϕ*_0_ < *θ* < *ϕ*_0_. Equation (12) has two possible solutions

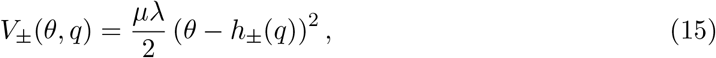

where +/− corresponds to the solution where the agent moves right/left. At 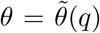, the two solutions meet. Since *V* is continuous, we have 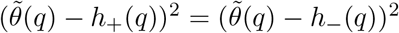 giving 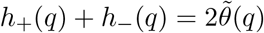. We re-write 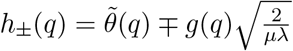(with *g*(*q*) > 0) to get

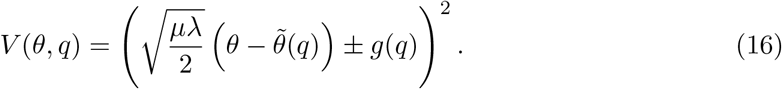

The above expression represents the cost of travel and connects the value at the boundaries; the two unknown functions 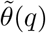 and g(q) can be found by matching the value functions at ±*ϕ*_0_. Note that *V* is positive as expected; if not, the agent would simply choose to stop and incur no control cost. From the equation for the optimal speed (10), for 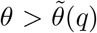, the agent moves to the right with speed 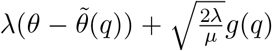 until it reaches *ϕ*_0_. The trajectory is therefore an exponential with exponent *λ*. By symmetry, we expect 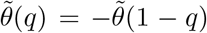 and *V* (*θ*, *q*) = *V* (−*θ*, 1 − *q*). In particular, 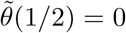, i.e., moving to the right or left are equally favorable if the agent is at *θ* = 0 and *q* = 1/2.

*θ* = *ϕ*_0_: Consider now the HJB equation at *θ* = *ϕ*_0_. By inspection, equation (13) has the particular solution q(1 + *λ*)^−1^. The general solution is then the particular solution plus a constant *C* times the homogeneous solution (1 − *q*)^1+*λ*^/*q*^*λ*^:

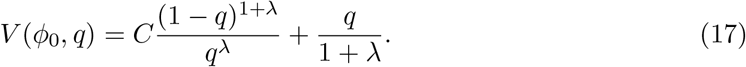

Note that the above expression satisfies the boundary condition *V* (*ϕ*_0_, 1) = (1 + *λ*)^−1^, consistent with the expectation that if the target is known to be at *ϕ*_0_ and the agent is at *ϕ*_0_, the expected discounted reward is 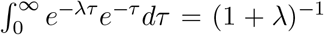. Equation (17) has an alternative, perhaps more intuitive, derivation. Recall the switching probability *q_s_* at which point the agent decides to leave *ϕ*_0_. The value at *V* (*ϕ*_0_, *q*) is the expected discounted reward of finding the target before the posterior q gets to *q_s_* plus the discounted value at *V* (*ϕ*_0_, *q_s_*) if the agent doesn’t find the target. Since 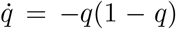, the sampling time for *q* to get to *q_s_* is 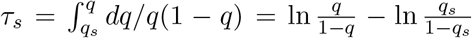. The expected discounted reward is therefore 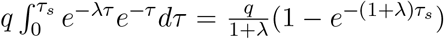. The probability of not finding the target in *τ*_*s*_ is 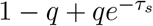. Combining these expressions we get

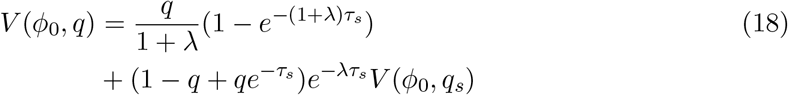

Plugging in the expression for *τ*_*s*_ yields (17). The equation holds for *q* ≥ *q_s_*, for any arbitrary *q_s_*. From the second derivation, it is clearer that the constant *C* in (17) contains all the dependencies on *q_s_*. Since the optimal switching point, 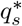, corresponds to the point at which the value is maximized, it is therefore the point at which *C*(*q_s_*) attains its maximum.

#### 2.3.1 Relationship with marginal value theory

Before proceeding to find 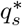, it is worth making a few clarifying points regarding the calculation of the optimal switching probability 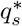 as the maximum of *C*(*q_s_*) and its relationship with marginal value theory.

In marginal value theory as applied to foraging, one computes the optimal switching point as the point at which the value of continuing to sample at a certain location equals the value of switching to another location. If the agent decides to continue sampling at 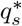, it would imply that (13) applies just below 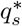 as well as above it, where 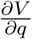 is instead the left derivative at 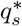. Continuity of *V* then implies that the left and right derivatives match at the optimal switching probability

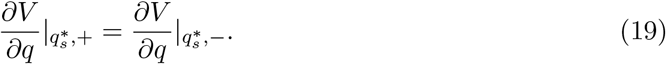

We now show that this condition is equivalent to maximizing *C*(*q_s_*) w.r.t *q_s_*. Call *f*_1_(q) = (1 − q)^1+*λ*^/*q*^*λ*^ and *f*_2_(*q*) = *q*/(1 + *λ*). From (17) we have for the right derivative 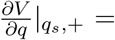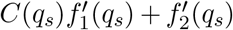. For *q* < *q_s_*, the agent immediately traverses from *ϕ*_0_ to −*ϕ*_0_. Using (16) at *ϕ*_0_ and −*ϕ*_0_ we get

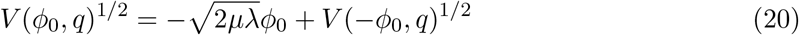

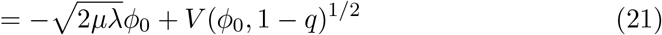

Equation (17) applies to *V* (*ϕ*_0_, 1 − *q*) since 1 − *q* > 1 − *q_s_* ≥ *q_s_*. Taking the derivative of (21) w.r.t *q* on both sides

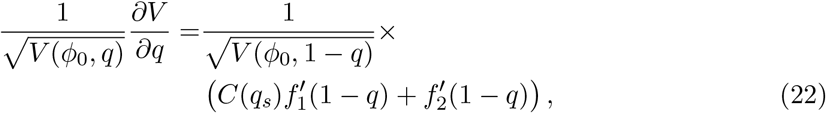

which yields the left derivative of *V* at *q_s_*. At the optimal switching point, equating the left and right derivatives at 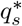, we get the condition for the optimal switching probability

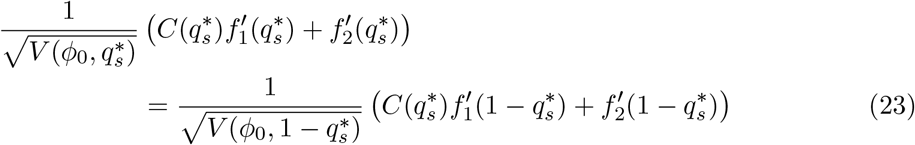

For an arbitrary switching point *q_s_*, using (16) at *q_s_*, we have

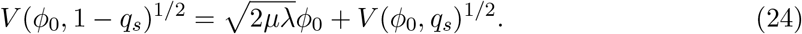

Plugging in (17) for the two value functions, taking the derivative w.r.t *q_s_* on both sides and imposing (23) at 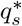, it is cumbersome but straightforward to show that dC(*q_s_*)/d*q_s_* = 0 at 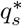.

#### 2.3.2 Optimizing for *q_s_*

Finally, to calculate the maximum of *C*(*q_s_*), we use (24) to first calculate *C*(*q_s_*) for any *q_s_*. Using (17) for *V* (*ϕ*_0_, *q_s_*) and *V* (*ϕ*_0_, 1 − *q_s_*) in (24), and solving for *C*(*q_s_*), we get

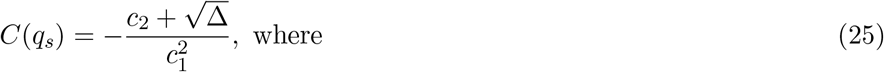

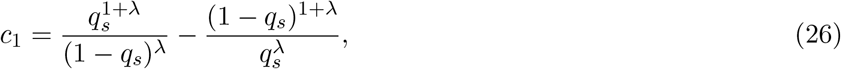

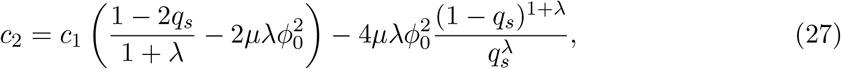

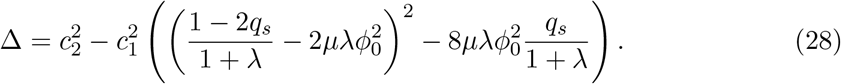

The value of 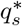 is then obtained using *dC*/*dq_s_* = 0 if there is a maximum for *q_s_* ≤ 1/2 or otherwise *q_s_* = 0. To get a sense of what the solution looks like, for *λ* ≪ 1 or when the cost of control is large (but below the point where it is no longer optimal to traverse), we expect 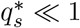. In this limit, we use (24) and match terms up to 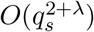 to get

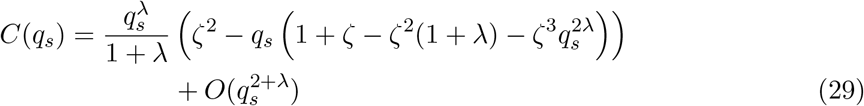

where 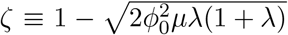. The above expression holds for *ζ* > 0. Equating *dC*/*dq_s_* to zero, we get

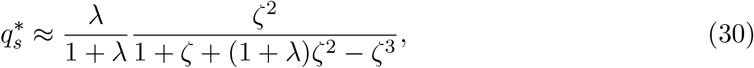

where we have used 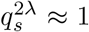 for *q_s_* ~ *λ* and λ ≪ 1. For *ζ* ≪ 1, that term can be ignored as its pre-factor is *ζ*^3^. For *ζ* ≤ 0, (24) cannot be satisfied, and we have 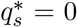 = 0. In Figure M2c, we verify the above approximation for 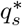.

### 2.4 Rate of information acquisition

An agent has a constant probability of finding the target per cycle. We compute the rate at which the probability of *not* detecting the target decays with time, which quantifies the rate of information acquired by the agent about the target. After 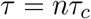 i.e., after *n* cycles of 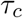 each, the probability of not detecting the target 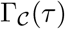 is

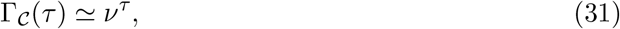

where 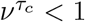 is the probability of not finding the target per cycle computed below. Denote 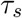 as the time spent sampling at *ϕ*_0_ (or −*ϕ*_0_) per cycle respectively. Since at *ϕ*_0_, the agent samples such that *q* goes from 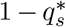 to 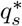, we have 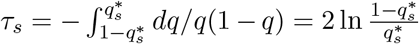. The probability of finding the target in this period is 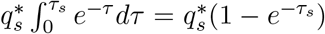. Therefore, we

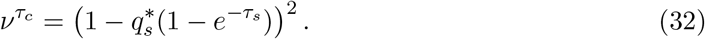

It remains to calculate 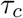. If *τ*_t_ is the time required for traversing from *ϕ*_0_ to −*ϕ*_0_ (or vice-versa), we have 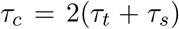. In (16) note that 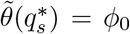 so that 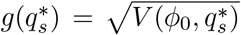 (which can be evaluated using (17)). The leftward speed when 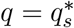 thus

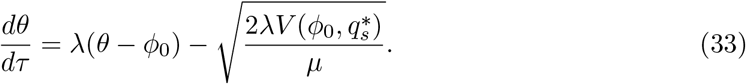

This is easily integrated to give

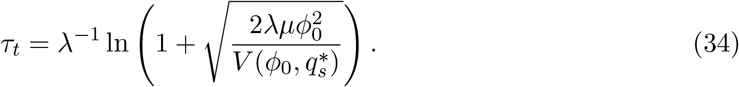

To summarize,

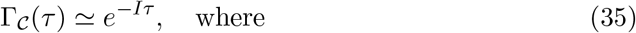

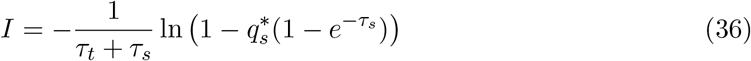

is interpreted as the rate of information acquisition. Reverting back to the original *t*-scale, we get

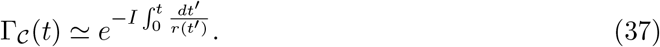

For the specific case of constant radial speed *v*, *r*(*t*) = *vt* + *r*_0_, the probability of not detecting decays as a power law,

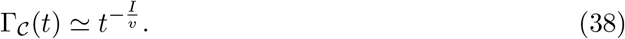

The mean time to find the target therefore diverges above a speed *v*\*/*aω* = *I*, which is set by a combination of the uncertainty *ϕ*_0_, the movement cost μ and the discount rate *λ*.

## 3 The Generalized Worm-like Chain (GWLC) ensemble of trails

Suppose *ϕ*_0_, *ϕ*_*L*_ are the headings at two points separated by distance L. We first compute the joint distribution *P*(*ϕ*_0_, *ϕ*_*L*_) for an Gaussian ensemble of trails with persistence in heading (WLC) and then introduce a new model with persistence in curvature, which we call the Generalized Worm-like Chain (GWLC) ensemble. While *P*(*ϕ*_0_, *ϕ*_*L*_) is the “interpolation” model, we will subsequently also calculate the forward model of how the trail’s heading and transverse position changes with distance from a fixed end.

Since *ϕ*_0_, *ϕ*_*L*_ are jointly normal, it is sufficient to compute their means and the covariance matrix. We denote *b*_1_, *b*_2_ for the diagonal and o*α*-diagonal elements respectively of the inverse covariance matrix. In the interpolation models, the agent makes contact with the trail at *x* = 0,*y* = 0 and *x* = *L*, *y* = 0. Small-angle approximation is used throughout; its validity is examined further below. 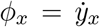 is the local heading (i.e., the tangent) and χ_*x*_ is the curvature.

### 3.1 Wormlike-chain

We begin by considering a simple wormlike-chain (WLC) model with no force, which corresponds to the model in the main text without curvature. The probability of a given path is

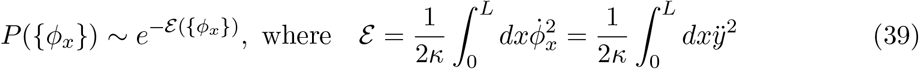

The Euler-Lagrange equation is *d*^4^*y*/*dx*^4^ = 0, so that the extremal path is a cubic polynomial, 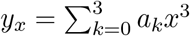. There are four parameters. Setting *y*_0_ = 0 and *y_L_* = 0, writing *a_k_*’s in terms of *ϕ*_*L*_, *ϕ*_0_ and plugging this in (39), we get the joint distribution

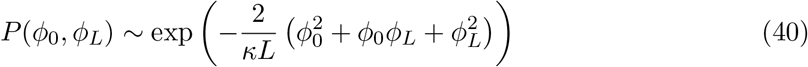

The inverse covariance matrix therefore has diagonal elements, *b*_1_ = 4/*κL* and off-diagonal elements, *b*_2_ = 2/*κL*. From inverting the matrix, the correlation is *ρ*_*L*_ = −*b*_2_/*b*_1_ and 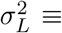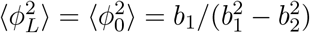, which gives

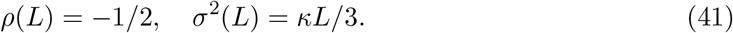

### 3.2 Fixed but unknown curvature

We now consider the case when the trail has a fixed but unknown curvature 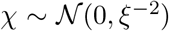, where *ξ* is the typical radius of curvature. Given *χ*, the WLC action is modified as

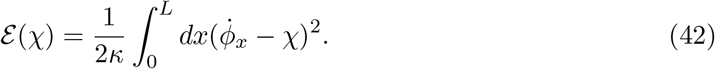

When the hidden variable *χ* is integrated out, we get

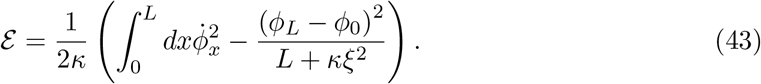

This looks much like the WLC case from before except for the extra boundary terms. These don’t affect the extremal path which is still a cubic polynomial. Plugging this in as before after setting *y*_0_, *y_L_* = 0, we get

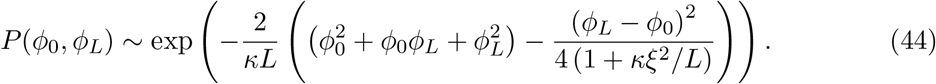

We can immediately see that when *κξ*^2^/*L* ≫ 1, this reduces to the WLC case. This is simply the case of (*κL*)^1/2^ ≫ *L*/*ξ* i.e., the curve diffuses much more than it curves. Call μ_*L*_ = *κξ*^2^/*L*.

Expanding the above equation, we have

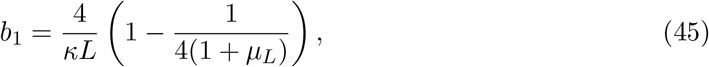

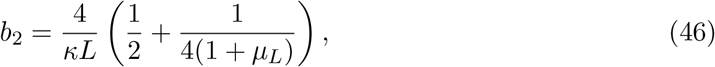

so that

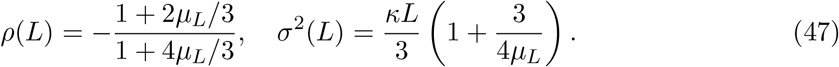

When *μ*_*L*_ ≫ 1, we recover *ρ*(*L*)≃ −1/2 and *σ*^2^(*L*) ≃ *κL*/3. On the other hand, when *μ*_*L*_ ≪ 1, we have

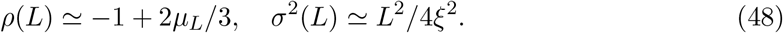

### 3.3 Varying curvature with a finite correlation length (GWLC)

In the previous section, we calculated the case of a fixed curvature, i.e., the curvature has infinite correlation length. Now, we assume the curvature is an Ornstein-Uhlenbeck process with correlation length *λ*. This corresponds to the full model considered in the main text. In this case, we have to integrate out the uncertainty over the initial curvature *χ*_0_ as well as the stochastic curvature field.

The full action including the *χ*_*x*_ curvature field and the stationary prior is

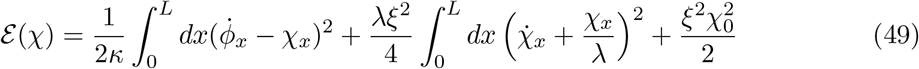

The prefactor *λξ*^2^/4 ensures the stationary distribution of *χ* is 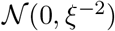. We integrate *χ*_*x*_ out using the Gaussian integral formula and obtain the e*α*ective action in *ϕ*_x_. To do this, we require the correlation matrix of the curvature process *χ*_*x*_, which we calculate below.

Let’s first calculate the correlation function for the curvature OU process without coupling it to the 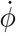 field, i.e., we ignore the first term in the action above. Let *χ*_0_, *χ*_1_,…, *χ*_*N*_ be the curvature at lengths 0, *dx*, 2*dx*,…, *Ndx*, where *dx* = *L*/*N* ≪ *λ*. The dynamics is defined by

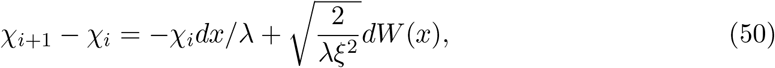

where 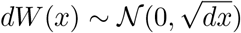. Then, the joint distribution of this set including the prior is

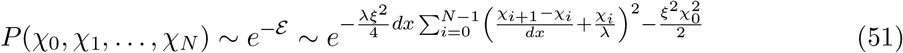

The cross term when the parenthesis is expanded is 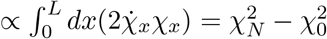. The action is

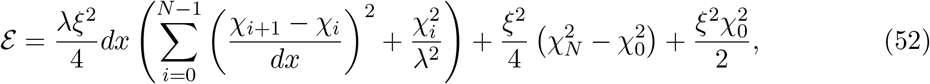

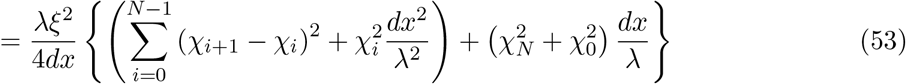

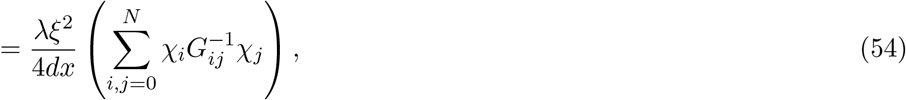

where *G* is the correlation matrix and *G*^−1^ is a tridiagonal matrix,

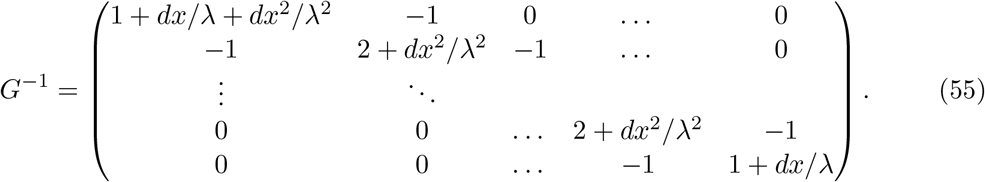

The inverse of a tridiagonal matrix can be calculated using a pair of recurrence equations [1]:

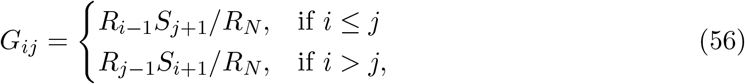

where *R* and *S* satisfy the recurrence relations and boundary conditions 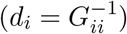

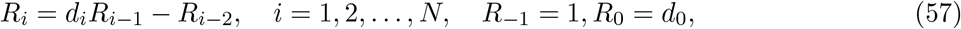

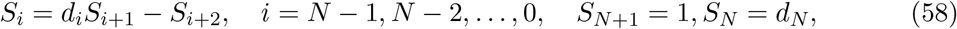

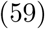

Since *d_i_* = 2 + *dx*^2^/*λ*^2^ for *i* ≠ 0,*N*, we can immediately see that in the continuous limit away from the boundaries

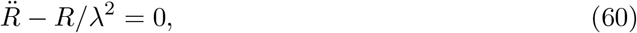

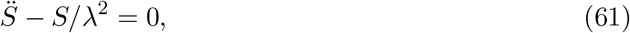

The difference between *R* and *S* is in the boundary conditions. We have *R*_−1_ = 1, *S*_*N*+1_ = 1, *R*_0_ − *R*_−1_ = *dx*/*λ* + *dx*^2^/*λ*^2^ and *S*_*N*+1_ − *S_N_* = −*dx*/*λ*, which implies in the continuum

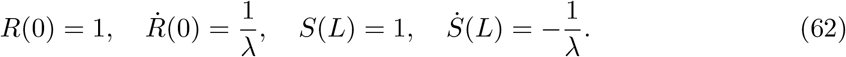

Solving for *R* and *S*, we get

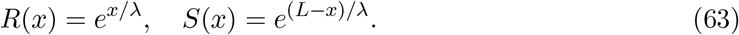

Finally, we should be careful about *R_N_* which appears at the denominator of the correlation matrix (56). We have *R_N_* = (1 +*dx*/*λ*)*R*_*N*−1_ − *R*_*N*−2_. *R*(*x*) is not continuous at the rightmost point. Call *R_N_* = *Z*(*L*), *R*_*N*−1_ = *R*(*L* − *dx*) and *R*_*N*−2_ = *R*(*L* − 2*dx*), we have as *dx*/*λ* → 0,

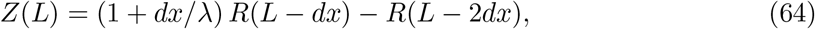

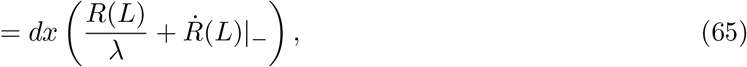

where 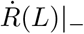 is the left derivative of *R* at *L*. We get *Z*(*L*) = (2*dx*/*λ*)*e*^*L*/*λ*^.

Plugging into (56), the correlation matrix in the continuous limit is

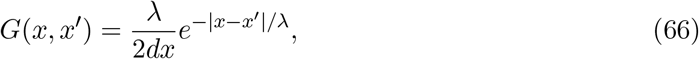

as expected for the correlation function of an OU process (note that we multiply by 2*dx*/*λ* in 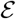).

Now, let’s get back to the full problem. The full action is

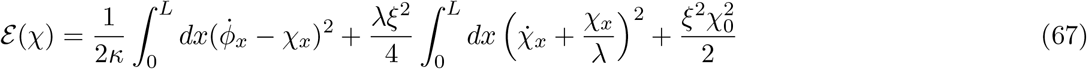

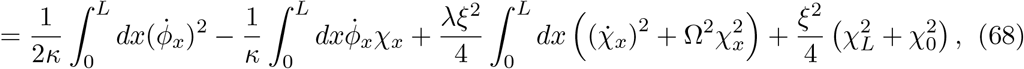

where the boundary terms are the same as above and we have defined *Ω*^2^ = *λ*^−2^ + *V*^−2^ and *V*^2^ = *κξ*^2^*λ*/2. We need the correlation matrix *G* of *χ* in this case.

However, notice now that even though the effective dynamics contain *Ω*^2^ instead of *λ*^−2^, the boundary conditions remain the same. That is, we should still get (62), but the solutions of (60) are now of the form *R*(*x*) = *Ae*^*Ωx*^ + *Be*^−*Ωx*^ (and similarly for *S*). We instead get

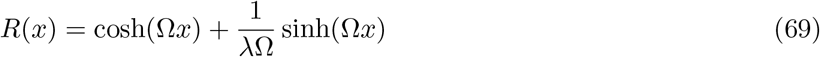

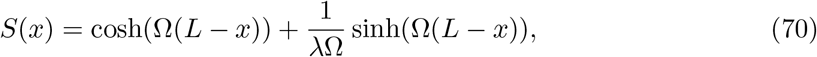

i.e., *A* = (1 + 1/*λΩ*)/2, *B* = (1 - 1/*λΩ*)/2. Since the boundary conditions are the same as above, we have for the normalization

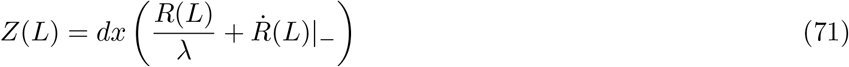

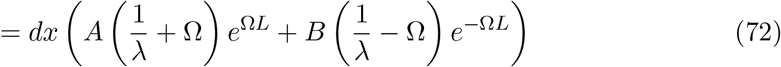

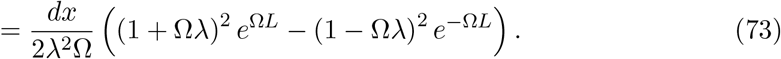

Once we multiply by the pre-factor, *λ*/2*dx*, the re-scaled correlation matrix, *G*(*x*, *x′*) is

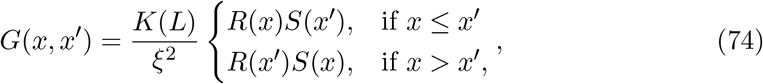

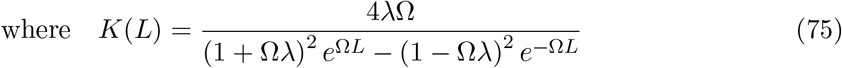

Using the Gaussian integral formula in (67), we get for the effective action

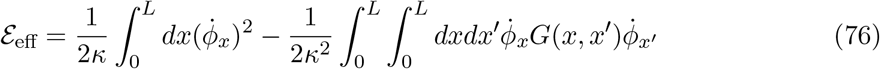

#### 3.3.1 Fixed curvature limit

We check if the case derived for fixed curvature is recovered when *λ* → ∞. Since 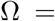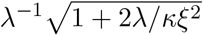 when *λ* is taken to be large, we get (*λΩ*)^2^ ≤ 2*λ*/*κξ*^2^. Also *ΩL* ≪1. From (75), in this limit,

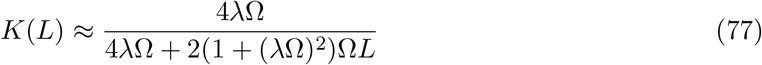

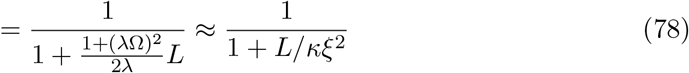

When *ΩL* ≪ 1, both *R*(*x*) and *S*(*x*) are 1. The correlation matrix is therefore

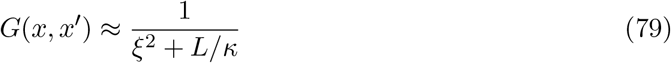

Plugging in (76), we get

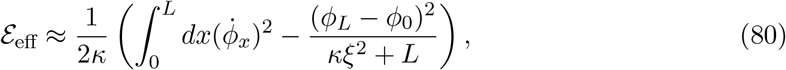

which is the one derived assuming fixed curvature.

#### 3.3.2 Extremal equation

With the correlation matrix in hand, we proceed to compute the extremal equation in *y_x_* satisfied by the effective action. From (76), the Euler-Lagrange equation is

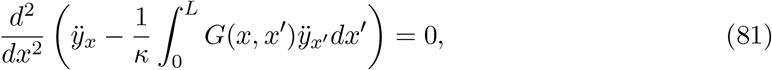

which gives

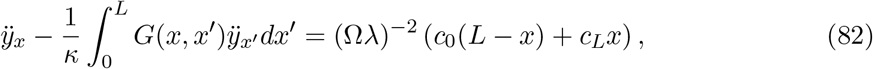

where the form of the r.h.s has been chosen for notational convenience which will be apparent later. *c*_0_ and *c_L_* are constants. The integral over *G* can be eliminated by differentiating w.r.t *x* twice. Define *f*(*x*) = *λR*(*x*). Then the correlation matrix is *G*(*x*, *x′*) = *K*(*L*)*f*(*x*)*f*(*L* − *x′*)/*λ*^2^*ξ*^2^ for *x* ≤ *x′* and *G*(*x*, *x*) = *K*(*L*)*f*(*L − x*)*f*(*x′*)/*λ*^2^*ξ*^2^ for *x* ≥ *x′*.

Differentiating once w.r.t x gives

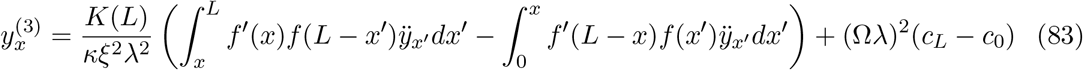

Differentiating once more, we can use

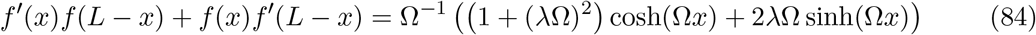

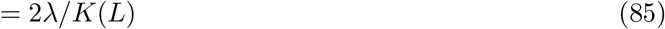

and *f″*(*x*) = *Ω*^2^*f*(*x*) and *f″*(*L* − *x*) = *Ω*^2^*f*(*L* − *x*) to get

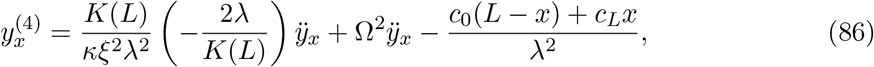

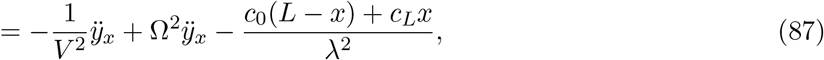

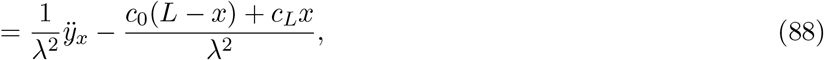

where we have used *Ω*^2^ = *λ*^−2^ + *V*^−2^ and (82) to get rid of the integral over *G*.

#### 3.3.3 The joint distribution of end-point angles

The general solution of the extremal equation (88) can be written as

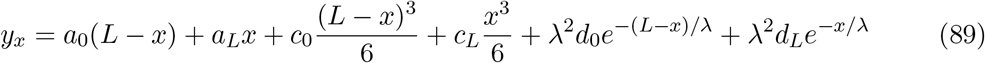

We’d like to get the coefficients in terms of *ϕ*_0_ and *ϕ*_L_ given *y*_0_, *y*_L_ = 0. The coefficients *d*_0_ and d_L_ are not independent of *c*_0_ and *c_L_* and are to be obtained by plugging in (89) into (82). For convenience, we work in terms of 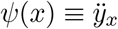. We have

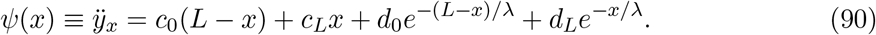

We can use integration by parts to compute the integral in (82). We have

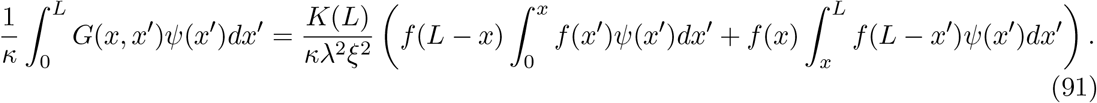

Write *ψ*(*x*) = *h*(*x*)+ *g*(*x*), where *h*(*x*) = *c*_0_(*L − x*)*R*+ *c_L_x* and *g*(*x*) has the exponential terms.

Then *h″*(*x*) = 0 and *g″*(*x*) = *g*(*x*)/)2. Let 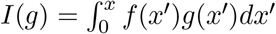. We have

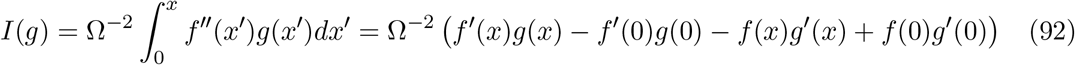

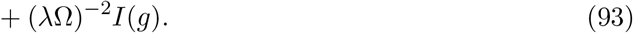

Sending the *I*(*g*) to the left side and using 1 − (*λΩ*)^−2^ = (*VΩ*)^−2^, we get

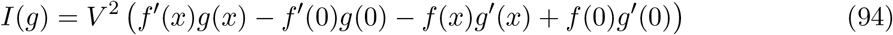

Similarly, define 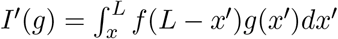.

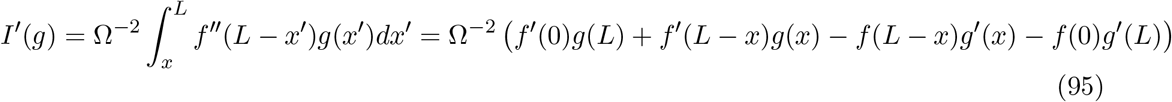

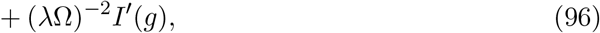

which gives

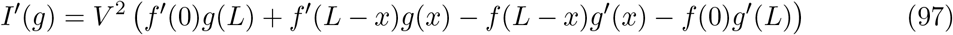

Eventually, after some simplifications we get

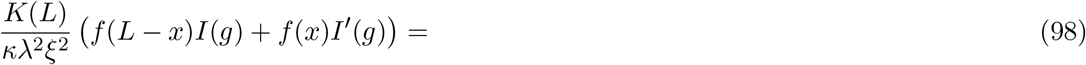

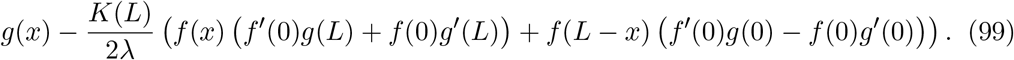

When we compute the same integrals for *h*(*x*) instead of *g*(*x*), since *h″*(*x*) = 0, when we integrate by parts we don’t get the *I*(*g*) as we did on the r.h.s of (93). We can replace *V*^2^ by *Ω*^−2^ and repeat the same calculation for 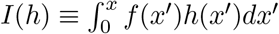 and *I′*(*h*) to get

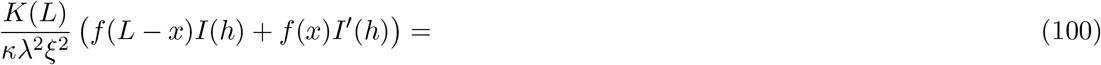

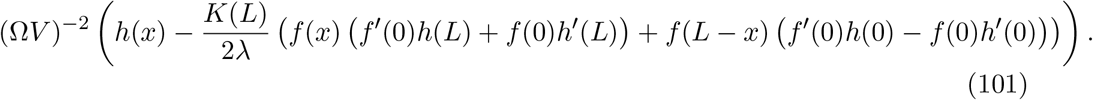

We subtract (99) and (101) from *ψ*(*x*) to evaluate the l.h.s of (82). The *g*(*x*) cancels out, whereas *h*(*x*)(1 − (*ΩV*)^−2^) = *h*(*x*)(*Ω*>)^−2^ cancels out with the r.h.s of (82). The leftover terms read

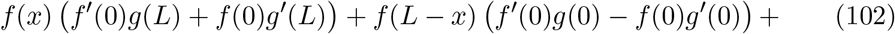

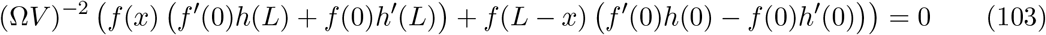

Define *m*(*x*) = *g*(*x*) + (*ΩV*)^−2^*h*(*x*). Using *f*(0) = *λ* and *f′*(0) = 1, we have

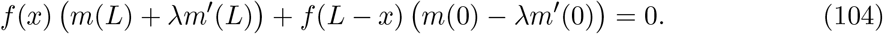

Since *f*(*x*) and *f*(*L* − *x*) are independent, we have *m*(*L*)+*λm′*(*L*) = 0 and *m*(0)− *λm′*(0) = 0, which we re-write as *m*(*L*)+ *m*(0) = *λ*(*m′*(0) − *m′*(*L*)) and *m*(0) − *m*(*L*) = *λ*(*m′*(0) + *m′*(L)). From the definition of *g*(*x*) and *h*(*x*), we have

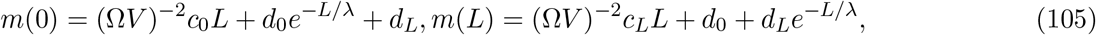

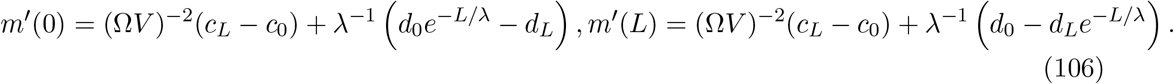

From the above two conditions *m*(*L*)+ *λm′*(*L*) = 0 and *m*(0) − *λm′*(0) = 0, it is easy to show that

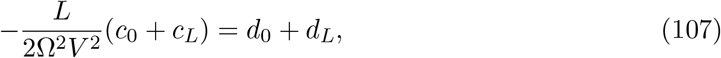

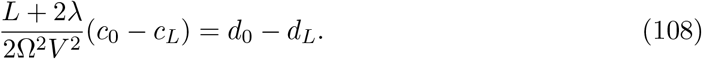

We want to write the effective action as a quadratic form in *ϕ*_0_ and *ϕ*_*L*_. Notice that the effective action can be written as

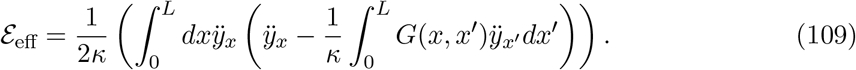

When we integrate by parts twice, two of the terms are zero, one because *y*_0_, *y_L_* = 0 and the other from the extremal equation (82). The leftover term is a boundary term

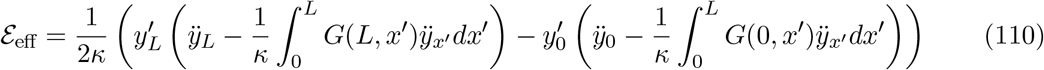

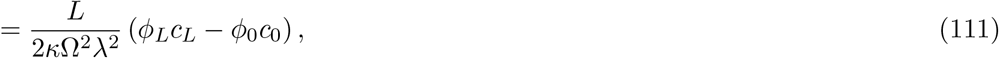

where we have used (82). It remains to express *c*_0_, *c_L_* in terms of *ϕ*_0_, *ϕ*_*L*_. The inverse covariance matrix should have equal diagonal elements and equal off-diagonal elements, and so we should have the form b_1_(*c_L_* + *c*_0_) = *ϕ*_*L*_ − *ϕ*_0_ and *b*_2_(*c_L_* − *c*_0_) = *ϕ*_*L*_ + *ϕ*_0_ for some *b*_1_, *b*_2_ which are to be found. From (89), we have

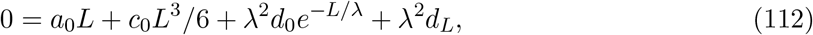

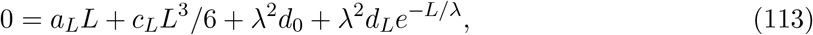

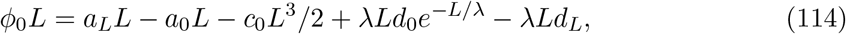

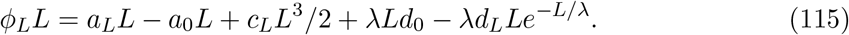

After some straightforward manipulations, we get

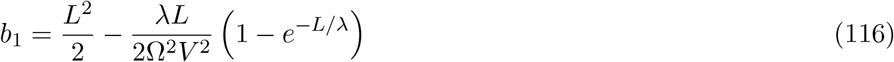

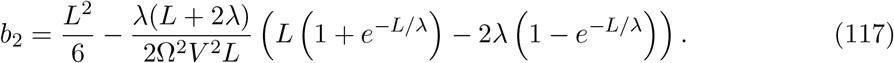

It can be checked that

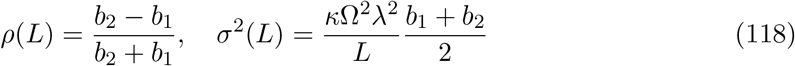

To recover the WLC and fixed curvature limits, we can expand *b*_1_ and *b*_2_ for small *L*/*λ* ≪ 1 to *O*(*L*^3^). We get

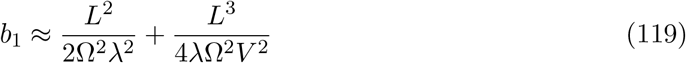

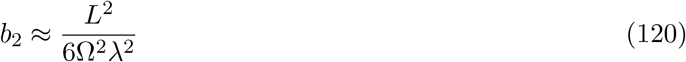

Plugging in (118), we recover *σ*^2^(*L*) ≤ *κL*/3+ *L*^2^/4*ξ*^2^.

#### 3.3.4 Numerical validation

Before comparing to numerics, we would like to examine the limits of validity of our original small-angle approximation, i.e., the conditions for 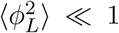 to hold. For simplicity, we consider the case when *κ* is small w.r.t the other length scales in the problem. Then, we can show that

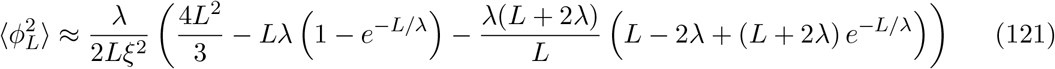

We assume *ξ* ≫ *λ* since otherwise the trail forms circles. When *L* ≫ *λ*, we can show diffusive scaling (also apparent in Figure M3c), 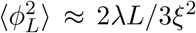. This expression is valid when 1 ≪ *L*/*λ* ≪ 3*ξ*^2^/2*λ*^2^. When *L* ≪ *λ*, (121) reduces to 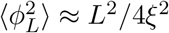 derived previously, which is always much less than one in that limit if *ξ* ≫ *λ*. Thus, the small-angle approximation is valid for *L*/*λ* ≪ 3*ξ*^2^/2*λ*^2^.

For numerical validation, we integrate over *s*, *dx*/*ds* = cos *θ*, *dy*/*ds* = sin *θ*, *dθ*/*ds* = *χ* and 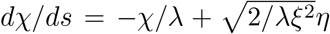, where *η* is white noise. The initial conditions are *x*(0) = *y*_0_ = *θ*(0) = 0 and 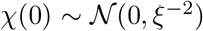. Rotating the *θ* axis so that initial and final *y* are zero, we have *ϕ*_0_ = −*y_L_*/*L* and *ϕ*_*L*_ = *θ*(*L*) − *y_L_*/*L*. We plot *σ* vs *L*, where 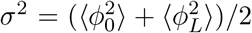. Results are shown in Figure M3e.

#### 3.3.5 Forward propagator

We exploit our previous calculation of the extremal path to compute the forward propagator, i.e., *P*(*ϕ*_*L*_, *y_L_*|*ϕ*_0_ = 0, *y*_0_ = 0) (if *ϕ*_0_ = 0, we can simply re-orient axes). In this case, the extremal path remains the same, but when we integrate the effective action (109) by parts we get a different expression:

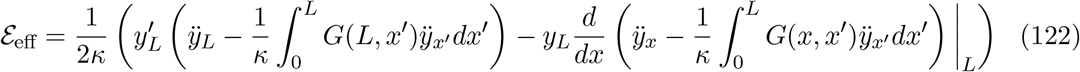

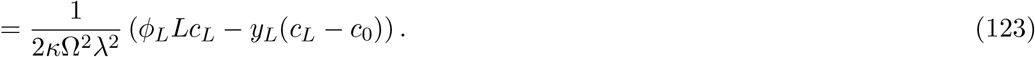

We need to express *c_L_* and *c*_0_ in terms of *ϕ*_*L*_ and *y_L_*. We have from (89)

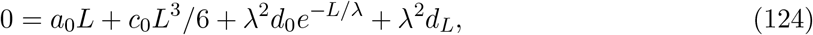

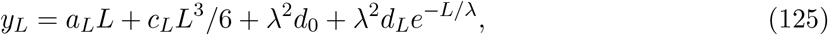

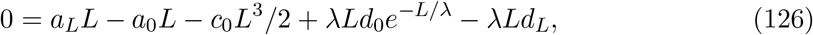

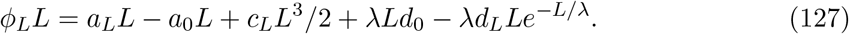

The relationship between *d*_0_, *d_L_* and *c*_0_, *c_L_* in (107) remains the same as before. From these equations, it is lengthy but straightforward to show that

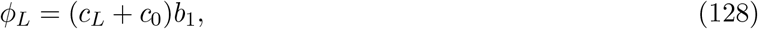

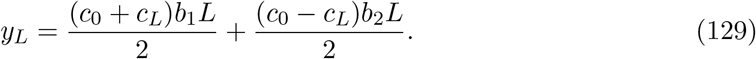

Solving for *c_L_* and *c*_0_ and plugging in to (123), we eventually get

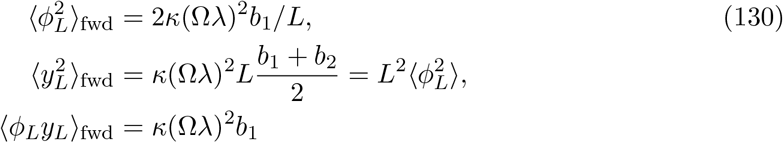

These expressions are numerically validated as shown in Figure M3f. Note also that the widening of the azimuthal position *y_L_*/*L* is given by 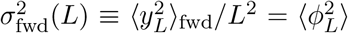, which is the variance of end-point angles for the interpolation model.

## 4 The non-detection probability during surge and cast

In this section, we analyze a sector search strategy where the agent casts rapidly and moves radially with constant speed. In contrast to the previous section, the full dynamics of trails is considered. The key quantity here is the non-detection probability of not detecting the trail at radial distance *L* after loss of contact, 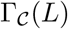, using which we can calculate mean times to detection and the probability of missing the trail. We compute the non-detection probability conditioned on initial path heading, 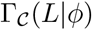. We have

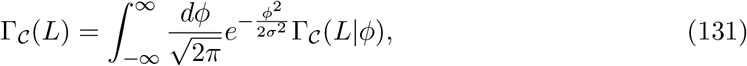

where *σ* is the uncertainty. We assume constant longitudinal speed *v*, transverse speed u, sampling rate *ω* and kernel size *a*. We also assume there is a duration, *t_r_*, after the most recent detection where the agent does not detect which represents the minimum delay after a sample (for e.g., the exhalation phase of a sniff). We expect *t_r_* ≈ *ω*^−1^. We set *t_r_* = 0 for the results presented in the main text and Methods. Throughout, we use the GWLC ensemble of trails (Section 3.3).

### 4.1 Non-detection probabilities in the cast phase

The probability that a particular trail path, {*y*(*x*)}, is not detected is the probability that the agent does not sample during the period when it is within detection range of the trail. The non-detection probability is thus *e*^−*ωT*^ ^({*y*(*x*)})^ where *T* ({*y*(*x*)}) is the total time of overlap between the agent’s detection range and the trail’s path. We assume the tangential speed *u*/*aω* ≫ 1 and the detection kernel has full-width a. Let the turning points be at the azimuthal envelope *σx*Θ_env_(*x*), where *x* is the distance along the most likely heading. During casting, the agent repeatedly traverses from −*σx*Θ_env_(*x*) to *σx*Θ_env_(*x*) (and vice versa) in time 2*σx*Θ_env_(*x*)/*u*. It is easy to show that if |*y*(*x*)| < *a*/2+*σx*Θ_env_(*x*), then the time spent within detection range of the trail during a single traversal is given by

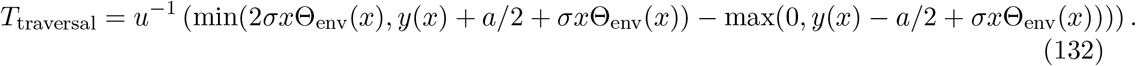

In the time it takes to travel *dx* in the longitudinal direction, the total number of traversals is 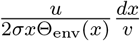. Therefore, the total time spent within detection range while the agent travels *dx* is

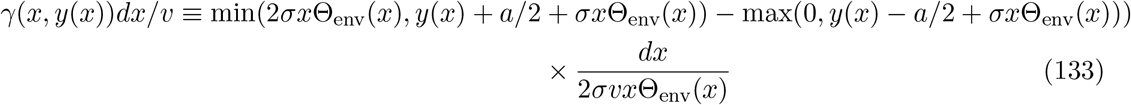

In this *u*/*aω* ≫ 1 limit, the detection probability is thus independent of *u*. To compute 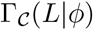, we need to integrate over all trail paths with *y*(0) = 0, *y′*(0) = *ϕ* up to length *L*.

That is

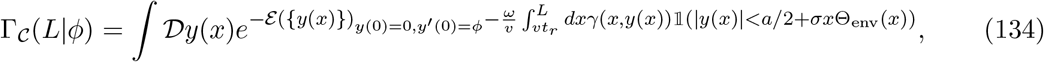

where 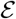 is the action defined by the specific model of the trail and the integral is over trails with *y*(0) = 0, *y′*(0) = *ϕ*. It is useful to rotate axes such that the paths *y*(*x*) have initial heading 0, i.e., equivalently,

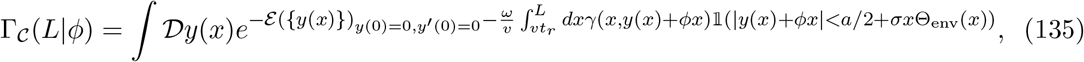

We now exploit the spline formulation from the previous sections. The simplification is that the extremal paths of the action have relatively simple forms like the one in eq. (89). Since the extremal paths are constrained only by the initial and final positions and headings, they have four parameters, two of which are fixed by *y*(0) = 0, *y′*(0) = 0. The action for the extremal case thus has two free parameters, viz., the final transverse position, *y_L_*, and final heading, *ϕ*_L_. From our previous results, the action is a quadratic form in *y_L_* and *ϕ*_*L*_, which have mean 0 and covariance 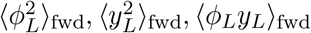 in (130). We will assume the diffusivity *κ* is negligible in the analysis hereafter. The results can be easily generalized to di*α*usive trails. Note that *ξ* ≫ *λ*, otherwise the trail forms circles. The path *y*(*x*) has the form (89), where the coefficients *a*_0_, *a_L_*, *c*_0_, *c_L_*, *d*_0_, *d_L_* can be written in terms of *y_L_*, *ϕ*_*L*_ using the relations derived previously. The expectation over paths in (135) is replaced with an expectation over *ϕ*_*L*_, *y_L_*:

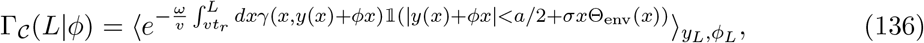

which is efficiently computed numerically by sampling (sampled *ϕ*_*L*_, *y_L_* can be re-used for all *ϕ*).

### 4.2 Geometric interpretation

When *v* is small, the first order contributions to the non-detection probability come from the path most likely to survive the casting search. That is, to first order we can approximate the integral (135) as

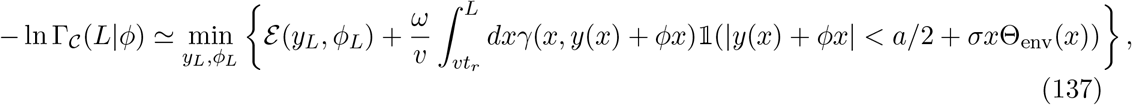

where 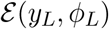, is the action. Note that the minimum is rather shallow, so the first order approximation is not a very good one (since the trails are flexible at the length scales of interest), but leads to some geometric intuition.

In Figure M4d we show examples of the path that minimizes the quantity in the parenthesis above. Intuitively, when *ϕ* is small, the minimizer path is ‘immersed’ in the rotated casting profile and should expend significant bending cost in order to escape the casting region. The minimizer is simply a straight line. In this case, 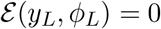 and all the *ϕ* within this range have approximately equal probability of being absorbed. As *ϕ* increases, the casting profile gets lower and there is a transition point, *ϕ*\* (which depends on L), where the minimizer path can escape the casting region without incurring too much bending cost. This is shown in the middle panel of Figure M4d. For *ϕ* > *ϕ*\*, the non-detection probability rapidly increases. At large angle, the casting profile no longer has a significant effect on the non-detection probability and the minimizer path begins to straighten out again.

The posterior distribution of the initial heading *P*(*ϕ*, *L*) is 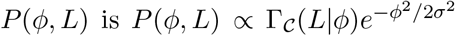, which can be numerically computed from (136). The prior is unimodal, which transitions to a trimodal shape at intermediate values of *L*. At large *L*, the posterior becomes bimodal. The locations of the modes at large *L* are computed below for small *v* and conical envelopes.

### 4.3 Conical casts

We analyze the case when Θ_env_(x) = Θ_0_ is a constant. In this case, 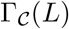 can be approximated and the optimal speed derived. The contribution to 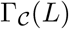 from trails outside the cone, |*ϕ*| > *σ*Θ_0_ is calculated further below. For trails directed inside the cone, their nondetection probability decreases independent of their initial heading until they escape the cone. The point at which they escape the cone is easily calculated. We give a heuristic argument (partly reproduced in the Methods) which captures the basic intuition and leads to a good approximation for 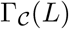.

The idea is that to first order, for *L*/*λ* ≪ 1, the trail paths have a single curvature scale *χ* ~ *ξ*^−1^. If |*ϕ*| < *σ*Θ_0_, the trails need to curve *σ*Θ_0_ − *ϕ* radians to leave the cone, which is at *L*\* ~ 2*ξ* (*σ*Θ_0_ − |*ϕ*|). The non-detection probability can be calculated from (136), but *γ* has a complicated dependence on x and y(x). To simplify, we assume for trails inside the cone *γ*(*x*, *y(x)*) ≈ *γ*(*x*, 0), which corresponds to the trail directed along the cone’s midline. Then, *γ*(*x*, *y(x)*) = 1 if *a*/2 > *σ*Θ_env_(*x*) and *γ*(*x*, *y*(*x*)) = *a*/2*σ*Θ_env_(*x*) otherwise. Define *β* ≡ *aω*/2*σv*Θ_0_. From (136), we get for |*ϕ*| < *σ*Θ_0_

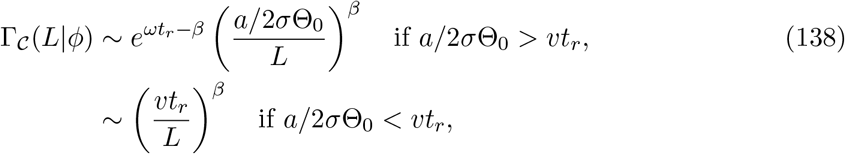

until it flattens out at some *L_in_* calculated below. For small *L* and Θ_0_ ≫ 1, 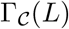 is determined by the trails directed within the cone and thus 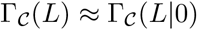 also decays as (138) until it flattens out. There are two possibilities for 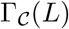 to flatten out, whichever occurs earlier: 1) a significant fraction of trails leave the cone, or 2) 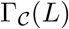 decays until it matches the probability of missing the trails that began outside the cone.

We estimate the length scale at which a significant fraction of trails leave as *L*_in_ ~ 2*ξσ*Θ_0_, when even the trails initially directed towards the center of the cone escape. This underestimates *L_in_* since the curvature of trails with large initial curvature (the ones that escape) may revert back to the mean. At *L* = *L_in_*, 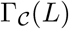 flattens out at

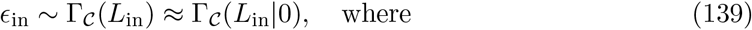

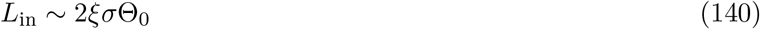

This fixed curvature estimate is valid when 2*ξσ*Θ_0_ ≪ *λ*. When *L* ≫ *λ*, the trails’ azimuth diffuses as *ϕ*^2^ ~ *λL*/*ξ*^2^, and thus it takes 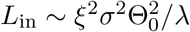 for most trails to diffuse *σ*Θ_0_ and out of the cone.

We now calculate the probability of missing the trails that began outside the cone, *ϵ*_out_, i.e., the trails with |*ϕ*| > *σ*Θ_0_. These trails leave the cone early before the casting phase 2*σ*Θ_0_*L* ≫ *a* begins. Since *L* is small, we can assume these are sti*α*, straight paths with angle *ϕ*. The distance at which they escape is *L*\* ~ *a*/2(|*ϕ*| − *σ*Θ_0_). Define *β′* ≡ *σv*/*aω*. The conditional non-detection probability is

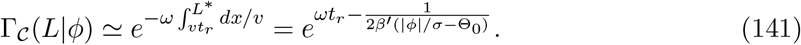

Then,

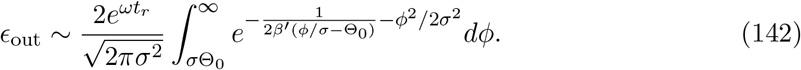

Transforming variables *ϕ* → *ϕ*/**σ** − Θ_0_ and expanding the prior term, we get

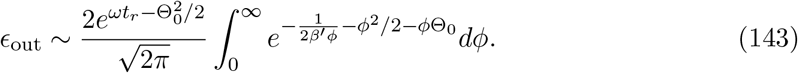

We calculate the above integral in the small *β*′ limit. This is because, as shown below, when *β′* ≪ 1, the detection rate of trails towards the center of the cone scales as *β′*^−1^, whereas for trails that begin outside the cone the detection rate scales as *β′*^−2/3^ or *β′*^−1/2^. Thus the small *β′* limit is the relevant one for *ϵ*_out_ i.e., when *ϵ*_out_ > *ϵ*_in_.

When *β′* ≪ 1, (143) can be approximated using Laplace’s method, but the maximum is hard to calculate analytically since its a cubic root. It is easy to show that depending on whether 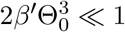 or 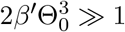, either *ϕ*^2^/2 or *ϕ*Θ_0_ respectively dominate at the maximum. For the two cases, we substitute *ϕ* = *β′*^−1/3^*ϕ′* or *ϕ* = *β′*^−1/2^*ϕ′* respectively to transform the moving maxima and use Laplace’s method to get up to constant prefactors

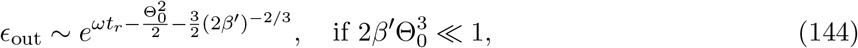

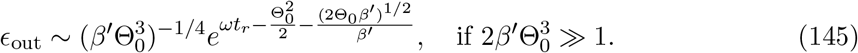

When *ϵ*_out_ > *ϵ*_in_, we can define a length scale, *L*_out_ when 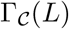 flattens out, which satisfies

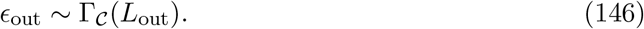

We can also estimate the initial heading of the trails that are most likely missed. When *ϵ*_out_ < *ϵ*_in_, this is the trail directed towards the center of the cone, *ϕ* ≈ 0, since the trails missed are the ones inside the cone and *ϕ* = 0 has the highest prior probability. When *ϵ*_out_ > *ϵ*_in_, this is located at the extrema in the integrals above, i.e., |*ϕ*|/**σ** = Θ_0_ + (2*β*′)^−1/3^ and 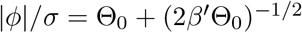 for 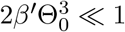 and 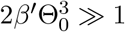 respectively.

In summary, we estimate if *a*/2**σ**Θ_0_ > *vt_r_*,

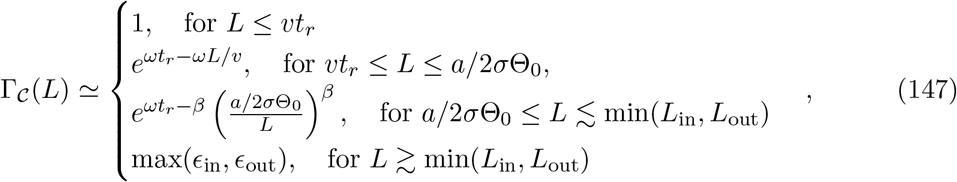

If *a*/2**σ**Θ_0_ < *vt_r_*, we have

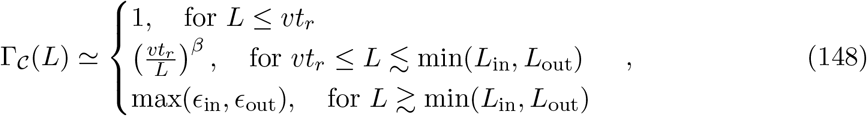

In Figure M5a we compare the prediction for 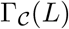 to the numerically obtained result using (136). Figure M5b shows the comparison between numerics and the prediction for 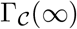 for a range of values of *β* and Θ_0_.

